# Core Clock Protein Subcellular Dynamics Coordinate Local and Global Circadian Control in Syncytia

**DOI:** 10.1101/2025.09.30.679467

**Authors:** Ziyan Wang, Bradley M. Bartholomai, Bin Wang, David Ritz, Daniel Schultz, Jennifer J. Loros, Jay C. Dunlap

**Affiliations:** Department of Molecular and Systems Biology, Geisel School of Medicine at Dartmouth, Hanover, NH, USA; Department of Microbiology and Immunology, Geisel School of Medicine at Dartmouth, Hanover, NH, USA; Department of Biochemistry and Cell Biology, Geisel School of Medicine at Dartmouth, Hanover, NH, USA

**Author notes:** Corresponding author: J. C. Dunlap.

## Abstract

Circadian rhythms are pervasive among eukaryotes, and the underlying clocks share a common regulatory architecture - a negative feedback loop. A wealth of genetic and biochemical data underpin current perceptions of circadian oscillators but aspects of their cell biology remain cryptic, especially in syncytial systems. We employed novel microfluidic systems and a blind mutant that retains circadian function to simultaneously track multiple clock components *in vivo* across circadian cycles, revealing remarkable subcellular and subnuclear dynamics of clock proteins and providing insights into spatiotemporal regulation in a multinucleated system. Despite heterogeneity of clock gene (*frq*) expression, we find robust cycles in FRQ nuclear localization among all nuclei and document free diffusion of multiple clock components among nuclei. Within nuclei, clock components form distinct, small, highly dynamic nuclear bodies that persist throughout the cycle, occasionally co-localizing for circadian regulatory functions. This rich context of *in vivo* spatiotemporal information illustrates how separate nuclear clocks ensure synchronous regulation of cellular activities across a macroscopic syncytium.

## INTRODUCTION

Circadian clocks are life’s internal timing device, providing evolutional benefits by helping an individual cell or organism anticipate and better prepare for daily environmental changes (Taton et al., 2020; Bass and Lazar, 2016; Jabbur et al., 2024; Larrondo and Canessa, 2019; Xu et al., 2022; Klarsfeld and Rouyer, 1998; Atamian et al., 2016). All eukaryotic circadian clocks are driven by transcription-translation negative feedback loops (TTFLs) that cycle with an intrinsic period of approximately 24 hours, and those of fungi and animals share a conserved regulatory architecture (Cox and Takahashi, 2019; Dunlap, 1999; Loros, 2020; Patke et al., 2020; Yuan et al., 2022). These feedback loops are self-sustaining yet can be reset by external cues, such as light and temperature, ensuring robust while adaptive rhythmicity.

*Neurospora crassa* is a well-established model for circadian biology and has contributed significantly to our understanding of eukaryotic clock mechanisms through extensive genetic and biochemical studies (Dunlap and Loros, 2017; Loros, 2020). The Positive Arm of the *Neurospora* TTFL is the White Collar Complex (WCC), a heterodimer of WC-1 and WC-2, which functions as both a transcriptional factor and a blue-light photoreceptor (Crosthwaite et al., 1997; He et al., 2002; Froehlich et al., 2002; Cheng et al., 2003); it shares sequence similarity with and is functionally equivalent to the CLOCK:BMAL1 heterodimer that is the Positive arm of mammalian clocks. At the subjective dawn, WCC drives the expression of *frq* transcripts, which encode the key Negative Arm component FRQ (Aronson et al., 1994a; b). Following translation, FRQ acts as a scaffold, functionally equivalent to the PER proteins in mammals, and assembles additional components, including FRH and CKI, to form the Negative Arm complex FFC (Dunlap and Loros, 2017). FRQ undergoes progressive phosphorylation, entering the nucleus and bringing CKI as part of the FFC to phosphorylate and thereby inhibit WCC activity and turn off *frq* transcription during the subjective day (Garceau et al., 1997; Luo et al., 1998; Cheng et al., 2005; Schafmeier et al., 2005; He et al., 2006; Baker et al., 2009; Tang et al., 2009; Shi et al., 2010; Hurley et al., 2013; Wang et al., 2019). This inhibition is released when FRQ becomes hyperphosphorylated at subjective night, releasing WCC to restart a new cycle (Lee et al., 2000; Denault et al., 2001; Baker et al., 2009). The core design of this feedback loop is conserved across fungi and animals (Cox and Takahashi, 2019; Greenwood and Locke, 2020; Loros, 2020; Liu and Bell-Pedersen, 2006; Dunlap, 1999; Patke et al., 2020), and although the specific proteins differ, these clock proteins share a common theme of bearing large regions of intrinsic disorder (Pelham et al., 2020; Querfurth et al., 2011; Hurley et al., 2013; Partch et al., 2005; Czarna et al., 2011; Michael et al., 2017; Dunlap and Loros, 2018; Liu et al., 2006).

*Neurospora* biology is of particular interest due to its syncytial organization (Mela et al., 2020; Jalihal and Gladfelter, 2025). In *Neurospora*, many nuclei share the same cytoplasm, with incomplete septa that partially compartmentalize the hyphae while allowing rapid exchange of proteins and even nuclei themselves between compartments (Freitag et al., 2004; Fischer, 1999; Bowman, 2025; Schmieder et al., 2019; Wang et al., 2023b). Syncytia play significant roles in development and function in various organisms from fungal hyphae to mammalian placenta and muscle fibers, and raise both challenges and opportunities for cellular coordination (Abmayr and Pavlath, 2012; Johannsen, 1941; Benirschke et al., 1998; Ejiri, 1983; Płachno and Świątek, 2011). In syncytia, including *Neurospora crassa*, nuclear autonomy is well characterized: individual nuclei can operate differently in cell-cycle progression, transcriptional activity, and potentially circadian timing (Pieuchot et al., 2015; Roper et al., 2013; Dundon et al., 2016; Zacchetti et al., 2018; Roberts and Gladfelter, 2015). This raises a fundamental question: in a common cytoplasmic environment, how is coherent rhythmic output achieved? Cell biological approaches have recently begun to address such questions in circadian systems (Xie et al., 2023; Yang et al., 2020; Cohen et al., 2014; Öllinger et al., 2014; Smyllie et al., 2016; Gabriel et al., 2021; Xiao et al., 2021; Koch et al., 2022; Smyllie et al., 2022, 2025; Lee et al., 2015). Live-cell imaging, in particular, allows direct visualization of clock protein dynamics in their native environment, offering spatiotemporal resolution unachievable with bulk biochemical assays although caveats have been suggested when fluorescently tagged proteins are overexpressed from non-native promoters (Xie et al., 2023). *In vivo* imaging of endogenous-level clock proteins has revealed new insights into intercellular communication and subnuclear organization of clock mechanisms in cellularized systems (Xiao et al., 2021; Yuan et al., 2022; Xie et al., 2023; Smyllie et al., 2025). In syncytial systems, where each nucleus could in principle operate an independent clock, cell biology approaches are especially essential to uncover how local interactions and molecular exchanges lead to circadian coordination. RNA imaging in *Neurospora* using single-molecule fluorescence in situ hybridization (smFISH) uncovered unsynchronized *frq* transcription between neighboring nuclei, and that the transcripts often cluster near the nucleus instead of being evenly spread throughout the cytoplasm (Bartholomai et al., 2022). This striking finding naturally leads to the question whether similar local heterogeneity also occurs at the protein level. However, previous attempts to visualize circadian protein dynamics in *Neurospora* have been limited by technical barriers, including light-induced clock-resetting by wavelengths needed to excite fluorophores and insufficient imaging toolboxes (Castro-Longoria et al., 2010; Schafmeier et al., 2008).

Here, we establish a long-term, high-resolution live-cell imaging platform for *Neurospora* by combining the blind *wc-1^ΔLOV^* allele (He et al., 2002; Wang et al., 2016) with customized microfluidic devices, enabling continuous observation for over two circadian cycles at high spaciotemporal resolution without light resetting. We employed fluorescently tagged proteins driven at wildtype levels by their own native promoters, and use of *wc-1^ΔLOV^* eliminated the constraint of using only long wavelengths for excitation (Castro-Longoria et al., 2010). Use of bright monomeric fluorophores from other spectral channels to visualize clock components at their endogenous levels *in vivo* (Wang et al., 2023b) thus allowed multiplex imaging to simultaneously examine the relationship between the Positive and the Negative Arms of circadian clock, as well as precisely distinguishing between the nuclei and cytoplasm using cellular markers. Using this system, we show that FRQ accumulates in nuclei and oscillates in synchrony across them, and that circadian clock proteins are shared among nuclei in support of coordinated timing. At the subnuclear level, both the Positive and the Negative Arms form dynamic nuclear bodies that persist throughout the cycle, with the oscillating interactions between them correlating with changes in the mobility of WCC nuclear bodies and therefore their functions. These findings reveal how the circadian clock is organized and coordinated in a syncytial context and highlight the remarkable subcellular and subnuclear dynamics of core clock proteins, providing new insights into temporal regulation in multinucleated systems.

## RESULTS

### Blind *wc-1^ΔLOV^* mutation and customized microfluidic devices facilitate continual microscopic imaging of live *Neurospora*

Dissection of the subcellular dynamics of circadian clock proteins in fungi requires the simultaneous resolution of three distinct problems - light quality, detection sensitivity, and rapid hyphal growth. The first problem is that light serves as the dominant circadian time cue for all organisms, rapidly resetting their circadian systems to subjective dawn; to follow a circadian cycle of clock protein subcellular location, light used for fluorescence excitation cannot be wavelengths that constantly activate the photoreceptor and reset the clock. Photoreceptor gene knockouts cannot be used because the principal fungal photoreceptor is WC-1, the same protein that is required for clock function in the dark (Crosthwaite et al., 1997). Previous research attempted to address this problem by using only red light excitation (561nm) to which WCC is less sensitive (Castro-Longoria et al., 2010); however, this not only sacrificed the sensitivity needed for precision visualization of lowly expressed clock proteins like FRQ and WC-1, but also eliminated the ability to multiplex (Wang et al., 2023b). To retain the use of high intensity blue-excitable fluorophores, we instead employed a “blind” allele of *wc-1* (*wc-1^ΔLOV^*) in which the coding sequence for the LOV domain is deleted so that the clock cannot respond to light but retains its ability to operate and be reset by temperature steps (He et al., 2002; Wang et al., 2016). Interestingly, *wc-1^ΔLOV^* still displays more robust conidiation under light compared to dark, suggesting activity of other non-clock photoreceptor(s) (Fig. S1A-B), so to make sure the loss of the LOV domain only affects the circadian light-sensing function but not other core clock functions, we assayed the clock in *wc-1^ΔLOV^* using a reporter that directly reports WCC activity, the *frq c-box* driving luciferase (Gooch et al., 2008). The strain has a normal compensated circadian clock across temperatures (Fig. 1A-B, Fig. S1C) and retains expression of core clock proteins (Fig. 1C) (Wang et al., 2016).

**Figure 1.**
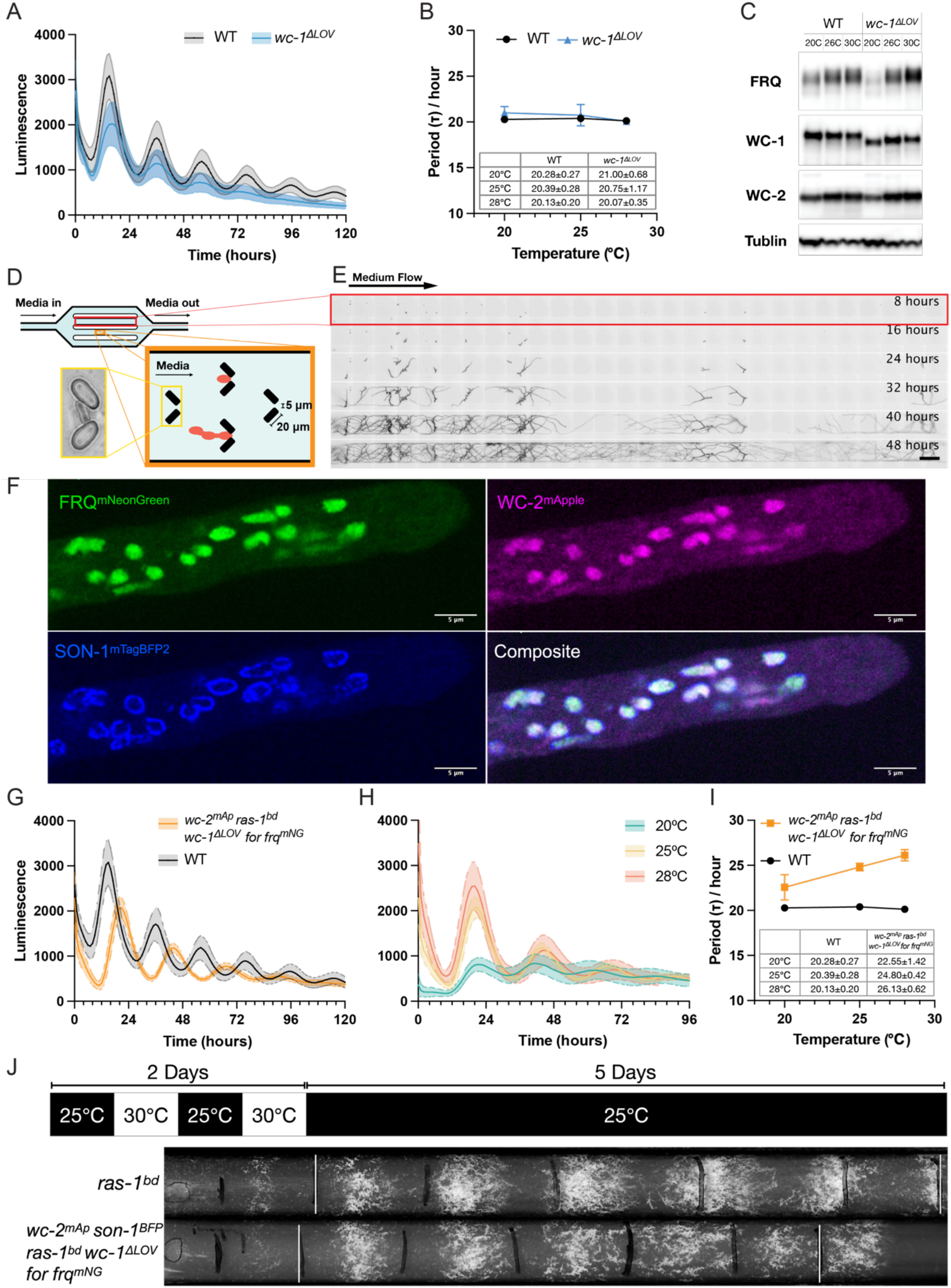
Blind *wc-1^ΔLOV^* allele mutation and customized microfluidic devices facilitate continual microscopic imaging of live *Neurospora*. (A) Luciferase assays of clock wildtype and *wc-1^ΔLOV^*strains at 25℃. Each curve represents the average of three biological triplicates each with 4 technical replicates, with the shade showing standard deviations. Here and in subsequent figures, the clock wildtype strain has the firefly luciferase gene under the control of the *frq* C-box inserted at the *his-3* locus (strain 661-4a, *his-3::frq_cbox_-luc*). (B) Periods calculated from luciferase assays of wildtype and *wc-1^ΔLOV^*strains at different temperatures following temperature entrainments. n=12 replicates for each condition. Error bars show standard deviations. (C) Western blot showing the expression levels of core clock components in wildtype and *wc-1^ΔLOV^* strains across temperature. (D) Schematic of the customized microfluidic device used to track *N. crassa* growth and circadian mechanics in real time. Conidia (orange in cartoon) are trapped in fixed locations and are kept in place by constant flow of fresh medium. The bottom left inset shows an actual example under light microscopy. (E) Example time lapse showing the growth of *wc-2^mApple^ son-1*^mTagBFP2^, *ras-1^bd^, wc-1^ΔLOV^ for frq^mNeonGreen^* strain (1981-2) in the microfluidic device. Images were taken in DAPI channel monitoring the fluorescent signal of nuclear envelope marker SON-1^mTagBFP2^; fresh medium flows from left to right. Scale bar = 200μm. (F) Middle focal plane images of the tri-color imaging strain (1981-2) grown on agarose pads under constant light, showing nuclear envelope (blue) and the localization of FRQ (green) and WC-2 (magenta). (G-H) Luciferase assays of the dual-color imaging strain *wc-2^mApple^ csr-1::frq_cbox_-luc, ras-1^bd^, wc-1^ΔLOV^ for frq^mNeonGreen^* (1981-1) (G) at 25℃ compared to wildtype or (H) across temperatures. Each curve represents the average of three biological triplicates each with 4 technical replicates, with the shade showing standard deviations. (I) Periods calculated from luciferase assays of the dual-color imaging strain (1981-1) at different temperatures after temperature entrainments compared to wildtype. n=12 replicates for each condition. Error bars show standard deviations. (J) Race tubes of wildtype and the tri-color imaging strain (1981-2), with white bars marking the beginning and end of 5-day growth under constant conditions, and black bars marking the growth front every 24 hours.

The third problem is that although we want to observe clock protein dynamics over the course of entire circadian cycles in the same cell, Neurospora grows extremely rapidly in 3-dimensions. To address this problem, we developed a microfluidic device with arrays of small barriers (Crane et al., 2014) to trap cells in a defined area, keeping strains both healthy and microscopically observable for extended periods (Fig. 1D). The height of the channel is 8 μm, ensuring a single layer of hyphal growth, the PDMS material is oxygen-permeable, and a controlled flow of fresh medium provides nutrients and removes wastes. *for* and *ras-1^bd^* mutations were crossed into *wc-1^ΔLOV^* to achieve manageable growth rate within the small chamber (Fig. 1E, Fig. S1D) (Belden et al., 2007a; Wang et al., 2024). Surprisingly, we noticed a strong non-chemotropic trend of against-flow growth possibly related to mechanical force sensing (Fig. S1E, Supplemental Video 1). Bird medium with polyacrylic acid was used to minimize hyphal fusion and branching for easier segmentation of single cells in imaging processing (Kelliher et al., 2020).

FRQ and WC-2 were tagged with bright monomeric fluorescent tags mNeonGreen and mApple, respectively, to track the subcellular behaviors of both the Negative arm and the Positive arm of the core clock (Fig. S2A) (Wang et al., 2023b) and transformed into their endogenous loci to ensure normal regulation, creating the blind slow-grow fluorescent strain *ras-1^bd^ for wc-1^ΔLOV^ frq^mNeonGreen^ wc-2^mApple^* (strain 1981). An ectopically expressed nuclear pore complex marker, fluorescent-tagged SON-1, was added to facilitate identification of nuclear boundaries (Wang et al., 2023b; Roca et al., 2010). In constant light, both FRQ and WC-2 concentrate in nuclei (Fig. 1F). The blind slow-grow fluorescent strain retains an intact circadian clock and output while displaying the modestly altered circadian period and temperature compensation profile that is typical when FRQ is tagged at its C-terminus as in *frq^mNeonGreen^* allele (Fig. 1G-J, Fig. S2B), consistent with previous reports fusing various reporters to FRQ (Larrondo et al., 2012; Castro-Longoria et al., 2010; Baker et al., 2009). Strains with other tagged/mutated alleles retain wildtype clock properties (Fig. S2C-E) (Belden et al., 2007a; Wang et al., 2024). Importantly, the expression of core clock proteins is comparable between wildtype and mutated strains (Fig. S2F-G). Therefore, we determined the core circadian clock behaviors in this genetic background are representative.

Altogether these tools allow extended time-lapse imaging under the microscope at high spaciotemporal resolution to unravel the subcellular dynamics of core clock proteins over circadian cycles.

### FRQ translocates to nuclei and oscillates in synchrony at the subcellular level among nuclei

To visualize operation of the circadian oscillator in real time-lapse under constant conditions we monitored the subcellular dynamics of the Negative Arm (FRQ) and Positive Arm (WC-2) at 1-hour resolution for > 48 hours in strains (1981-2/3) bearing varied color nuclear markers (Supplemental Video 2). Cells were initially reset to the beginning of subjective day (Circadian Time (CT) 0) by temperature entrainment and kept in constant conditions. As expected, nuclear FRQ peaks roughly 10 hours after subjective dawn and oscillates with peaks and troughs roughly 24 hours apart (Fig. 2A-B). The changes in nuclear WC-2 level are mild except for the dramatic drop within the first 12 hours post entrainment possibly associated with temperature changes (Fig. 2A&C). The oscillation in FRQ and negligible oscillation of nuclear WC-2 are consistent with biochemical data (Garceau et al., 1997; Denault et al., 2001). Total hyphal FRQ quantified as mean fluorescent intensity was complicated, especially in the second cycle, by noise arising from the accumulation of auto-fluorescent vesicles in some cells during development (Fig. S3A-C).

**Figure 2.**
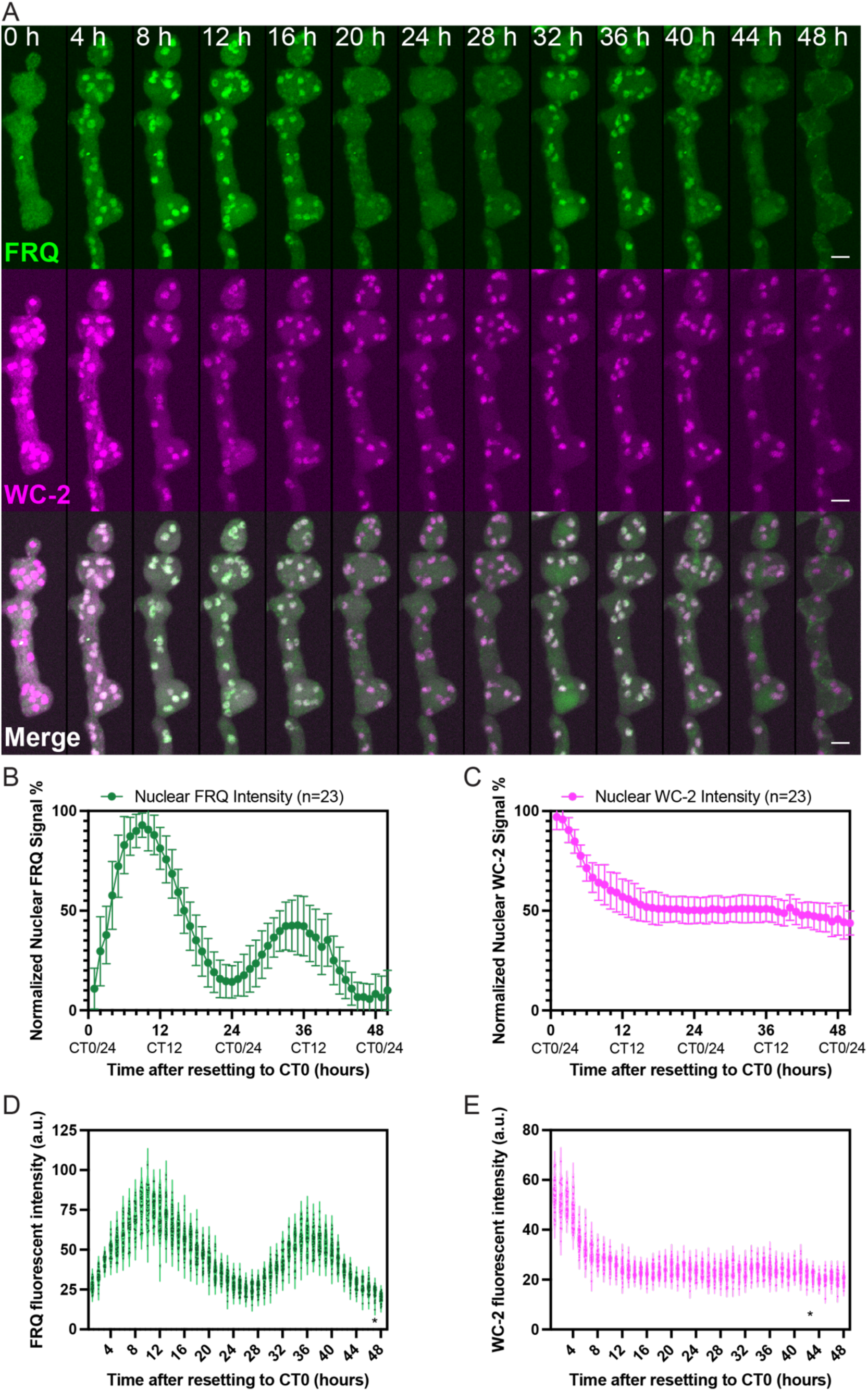
FRQ translocates to nuclei and oscillates in synchrony at the subcellular level among nuclei. (A) Localization of FRQ (green) and WC-2 (magenta) across circadian cycles. Strain 1981-2 grown in a customed microfluidic device (Fig.1D) was imaged every hour. Shown are maximal z-projections of z-stacks acquired from the same cell over time. Scale bar = 5μm. (B) Average fluorescent intensities of nuclear FRQ from n=23 cells. The maximal and minimal signal for each cell is set as 100% and 0% for normalization. (C) Average fluorescent intensities of nuclear WC-2 from n=23 cells. The maximal signal for each cell is set as 100% for normalization. All error bars show one standard deviation. (D-E) Distributions of (D) FRQ or (E) WC-2 signal in each nucleus over time for the example cell in (A). Each dot represents the mean intensity of one nucleus. * Indicates that the mean nuclear fluorescent intensity at the corresponding time point didn’t pass the D’Agostino-Pearson normality test (P ≤ 0.05).

An unexpected result is that FRQ visually concentrates in nuclei to some extent throughout the day (Fig. 2A), whereas most prior subcellular fractionation experiments had suggested that FRQ is mostly cytoplasmic (Luo et al., 1998; Hong et al., 2008; Diernfellner et al., 2009; Cha et al., 2011). However, when taking volume of subcellular compartments into consideration, the ratio of nuclear FRQ to total FRQ oscillates around 10%, similar to what was concluded from fractionation experiments (Fig. S3D). In conclusion, FRQ is always more concentrated in nuclei although derived from only a small portion of FRQ molecules.

Nuclear FRQ concentrations across individual nuclei are generally homogeneous, consistently adhering to a normal distribution without distinct subpopulations (Fig. 2D, Fig. S3E). This observation is striking considering that previous results showed *frq* transcripts are heterogeneously transcribed and distributed (Bartholomai et al., 2022). A similar homogeneous distribution pattern is seen for WC-2 proteins as well (Fig. 2E, Fig. S3F). Taken together, these results imply that protein distribution or sharing almost certainly happens in the shared cytoplasm, preventing any nucleus from drifting off on its own schedule.

### Local sharing of clock components facilitates circadian control

To dig deeper into how all nuclei communicate to stay temporally synchronized, we designed experiments utilizing *Neurospora*’s ability to form heterokaryons in which nuclei with different genotypes share the same cytoplasm (Roper et al., 2013). 2 genotypes were used to investigate FRQ, one with a wildtype circadian system but lacking the *frq_cbox_-luc* reporter described above that directly reports the WCC activity in the core clock, the other with the *frq_cbox_-luc* reporter but lacking the actual *frq* gene (Fig. 3A). Homokaryotic strains with only this second genotype lack FRQ so the feedback loop cannot close and WCC is never inhibited; they exhibit elevated arrhythmic luciferase activity. However, heterokaryons bearing these two nuclear types in the same cytoplasm regain normal circadian oscillations in luciferase signal (Fig. 3B). This means “clockless” nuclei with the *frq_cbox_-luc* reporter are receiving FRQ proteins encoded by the wildtype nuclei to inhibit WCC at the appropriate times. Circadian output, represented by oscillations in *clock-controlled gene-1* (*ccg-1)* promoter-driven luciferase signals, can be similarly restored with oscillation amplitude being dose-dependent to amount of luciferase reporters (Fig. S4A) (Wang et al., 2023b). Analogous experiments with WC-1 or WC-2 confirm that components of the Positive Arm of clock can also be shared among nuclei (Fig. 3C-D, Fig. S4B-C). In all cases the period of clock is remarkably insensitive to FRQ dosage reflecting robustness of the feedback loop (Fig. 3B, Fig. S4D).

**Figure 3.**
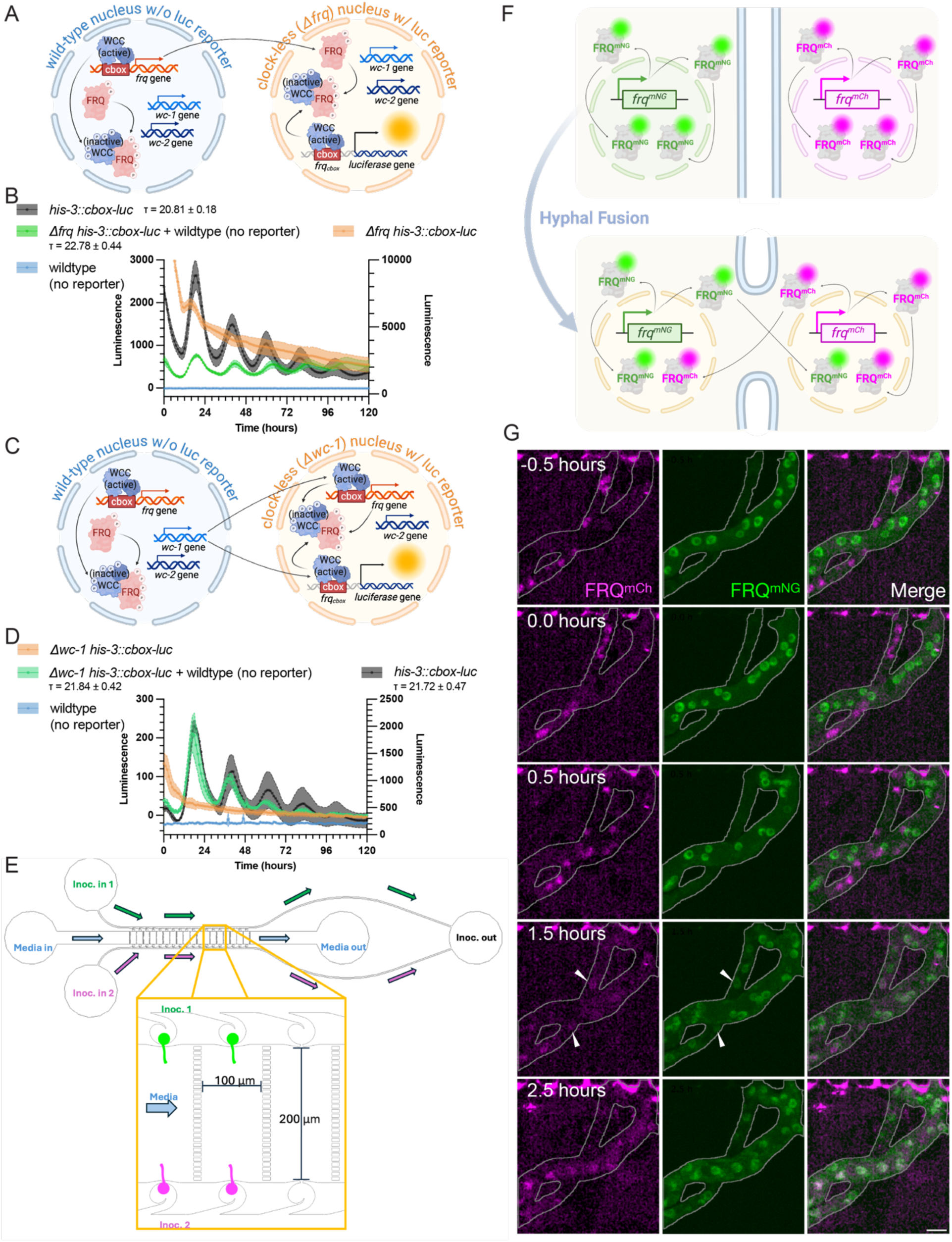
Local sharing of clock components facilitates circadian control. (A) Schematic showing the model for FRQ local sharing between nuclei in heterokaryons of clock-less reporter *Δfrq his-3::cbox-luc* and clock-competent reporterless wildtype. (B) Luc traces of *Δfrq his-3::cbox-luc*, wildtype strains, their heterokaryon, and the positive control strain *his-3::cbox-luc*. The scale for *Δfrq his-3::cbox-luc* is plotted on the right y axis, and the scale for the other 3 traces is plotted on the left y axis, as indicated above the axes. (C) Schematic showing the model for WC-1 local sharing in heterokaryon of *Δwc-1 his-3::cbox-luc* and wildtype. (D) Luc traces of *Δwc-1 his-3::cbox-luc*, wildtype strains, their heterokaryon, and the positive control strain *his-3::cbox-luc*. The scale for *his-3::cbox-luc* is plotted on the right y axis, and the scale for the other 3 traces is plotted on the left y axis, as indicated above the axes. In (B) and (D), each curve represents the average of 4 technical replicates with the shade showing one standard deviation. The periods (τ) of rhythmic traces are marked on the figure. For reference, plots with all traces, as well as another set of biological replicates, plotted on the same scale are included in Fig. S4E. (E) Schematic of the customized microfluidic device used to monitor local protein sharing in real time. (F) Schematic showing local sharing of FRQ variants after hyphal fusion. (G) Time-lapse imaging of FRQ^mNG^ and FRQ^mCh^ after fusion between *csr-1:: frq^mCh^; Δfrq* and *frq^mNG^* filaments. Scale bar = 5 μm. White arrowheads indicate representative nuclei with both FRQ^mNG^ and FRQ^mCh^.

To visualize this sharing behavior and reveal its timescale we developed another microfluidic assay in which cell fusion can be followed in real-time (Fig. 3E). We employed two strains, each expressing one FRQ variant driven by its endogenous promoter but marked by a distinct fluorescent tag (mNeonGreen or mCherry) (Fig. S4F-G). After inoculation of differing strains in opposite sides of the device, they grow against the medium flow towards the main chamber in the center, meet, and fuse with each other (Fig. 3E-F). Natural fusion events between hyphae can often be observed starting around 16 hours post inoculation (Supplemental Video 3). With local sharing of FRQ proteins, we expect to see gradual redistribution of FRQ^mNG^ and FRQ^mCh^ between nuclei regardless of their origins after hyphal fusion (Fig. 3F). Indeed, nuclei containing both FRQ variants start to appear within 1-2 hours after fusion and a new equilibrium can be reached soon thereafter (2-3 hours after fusion) (Fig. 3G). Heterokaryons of these 2 genotypes have pan-nuclear distribution of both FRQ variants at an intermediate concentration compared to individual homokaryons. This outcome resembles the equilibrium achieved after hyphal fusion, where the total nuclear FRQ level is comparable to the endogenous level, rather than being equivalent to expressing two copies of FRQ (Fig. 3G, Fig. S4G). Fast local sharing presumably facilitates the synchronization of all nuclei at the protein level, allowing them to function cohesively as a harmonious unit in driving circadian behaviors.

Overall, these experiments confirm that local protein sharing is a real phenomenon in circadian clock regulation and a major contributor to circadian synchronization in the syncytial tissue.

### Core clock proteins form highly dynamic nuclear bodies across circadian cycles

The subnuclear distributions of both Positive and Negative elements were plainly spatially heterogenous (Fig. 1F), so we investigated this phenomenon at higher spatiotemporal resolution by applying Spinning Disk Super Resolution by Optical Pixel Reassignment (SoRa) imaging.

Built upon a spinning disk confocal system, SoRa achieves an approximately 1.4x improvement beyond the optical limit in XY resolution through optical configuration, without requiring computationally intensive post-processing. Importantly, SoRa captures images at speeds comparable to a standard spinning disk confocal, making it suitable for imaging the highly dynamic behavior we see in live cells, a key advantage over many other super-resolution microscopes (Azuma and Kei, 2015). Using SoRa, we confirmed the presence of subnuclear FRQ foci, characteristic of phase-separated IDPs (Tatavosian et al., 2019; Caragliano et al., 2022; Tariq et al., 2024), that are similar in size and dynamics to nuclear microbodies of PER2, a Negative element in the mammalian clock (Xie et al., 2023). Indeed, both FRQ and WCC form highly dynamic nuclear bodies that do not remain stationary for more than 200ms (Fig. 4A, Supplemental Video 4). These ultrafast movements contribute to the blurry appearance within a single frame (100ms exposure). Sharper appearance of spots can be achieved with shorter exposure times but at the expense of volumetric information and signal-to-noise ratio (SNR) (Fig. S5A). Fusion or fission of FRQ nuclear bodies can be captured occasionally, consistent with *in vivo* liquid-like properties of nuclear FRQ (Fig. 4A-B).

**Figure 4.**
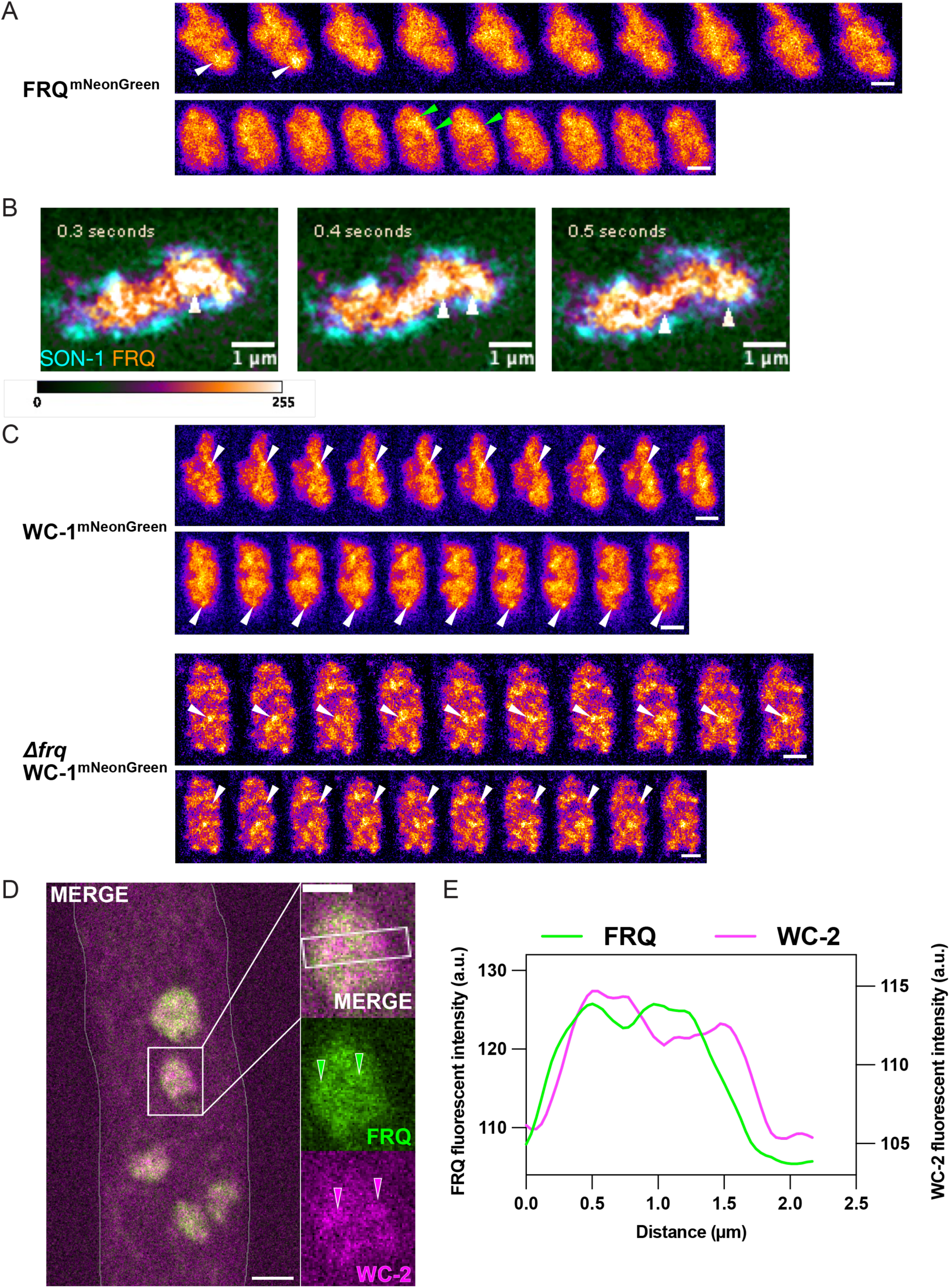
Core clock proteins independently form highly dynamic particles inside nuclei. (A-C) Timelapse images with 100ms exposure times without delays acquired on the SoRa system. All samples were cultured in constant light. (A) Two timelapses of FRQ^mNG^, each showing one nucleus. White arrowheads indicate a quasi-stationary nuclear body. Green arrowheads indicate a possible fusion event. Scale bar = 1μm. (B) Timelapse of FRQ^mNG^ showing an example fission event, indicated by white arrowheads. The “fire” look-up table at the bottom is for FRQ^mNG^. SON-1^mTagBFP2^ (cyan) marks the nuclear boundary. (C) Timelapses of WC-1^mNG^ in wildtype and *Δfrq* background. White arrowheads indicate one example of a stationary nuclear body in each timelapse. Scale bar = 1 μm. (D) Dual-camera images of FRQ^mNG^ and WC-2^mAp^ acquired simultaneously. Scale bar = 2 μm in the zoom-out (left). Scale bar = 1 μm in the inset (right). Green or magenta arrowheads indicate corresponding FRQ or WC-2 nuclear bodies along the white line shown in the merged inset. (E) Profile plot of fluorescent intensities in both channels along the white box in the inset in (D).

To facilitate a direct comparison between FRQ and WCC localization, we tagged the Positive element WC-1 (as a proxy for WCC) with mNeonGreen at its C-terminus inserted into its endogenous locus to ensure normal regulation and function (Fig. S5B) as well as ideal SNR and photostability for SoRa imaging. WC-1 nuclear bodies are significantly more stationary compared to FRQ while remaining heterogeneous and dynamic (Supplemental Video 5); some WC-1 nuclear bodies can be immobile for more than one second (Fig. 4C). Interestingly, while FRQ and WCC must function together in the clock, the existence of WC-1 nuclear bodies is an intrinsic property independent of FRQ as similar nuclear bodies are seen in *Δfrq* strains (Fig. 4C, Fig. S5C). Consistent with this, FRQ and WC-2 (as a proxy for WCC) nuclear bodies don’t always colocalize (Fig. 4D-E). The fast dynamics of intrinsically disordered clock proteins is not a universal feature for *Neurospora* nuclear proteins for histone H1 is immobile as expected (Fig. S5D, Supplemental Video 5).

As circadian rhythms are characterized by pronounced cycles in FRQ abundance and FRQ-WCC interaction, we had anticipated corresponding cycles in nuclear microbodies. However, surprisingly, time courses of *frq^mNG^* and *wc-1^mNG^* strains at 4-hour resolution show that nuclear bodies of both FRQ and WC-1 persist across circadian cycles, even when FRQ is at its trough level (Fig. 5A, Fig. S5E). To enable more precise volumetric characterization of nuclear bodies, as opposed to estimations derived from single optical sections of globular nuclei, we performed deconvolution on the 3D time-course data acquired at 1-hour resolution for downstream analysis (Fig. S5F). Similar to the single focal plane SoRa data (Fig. 5A), nuclear bodies of clock proteins can be observed at all times without exception (Fig. 5B). Nuclear bodies were identified from 3D reconstituted data using uniform parameters for all timepoints. Remarkably, our results indicate that the circadian oscillation in FRQ levels is not a reflection of a trivial change in the number of FRQ nuclear bodies within each nucleus, but rather the oscillation in the brightness of individual nuclear bodies (Fig. 5C-D, Fig. S5G). This change could be driven by the changing modifications and resulting physical properties of FRQ over circadian time as, in vitro, lower amounts of new non-phosphorylated FRQ are needed to initiate LLPS as compared to older phosphorylated FRQ (Baker et al., 2009; Tariq et al., 2024). The existence of similar numbers of FRQ nuclear bodies during the activation phase implies additional possible roles of nuclear FRQ as processing “hubs”, beyond inhibiting WCC (Pelham et al., 2023). For the positive element WC-2, as a proxy for WCC, alterations in both the brightness and the average number of nuclear bodies are minor (Fig. 5E-F, Fig. S5H).

**Figure 5.**
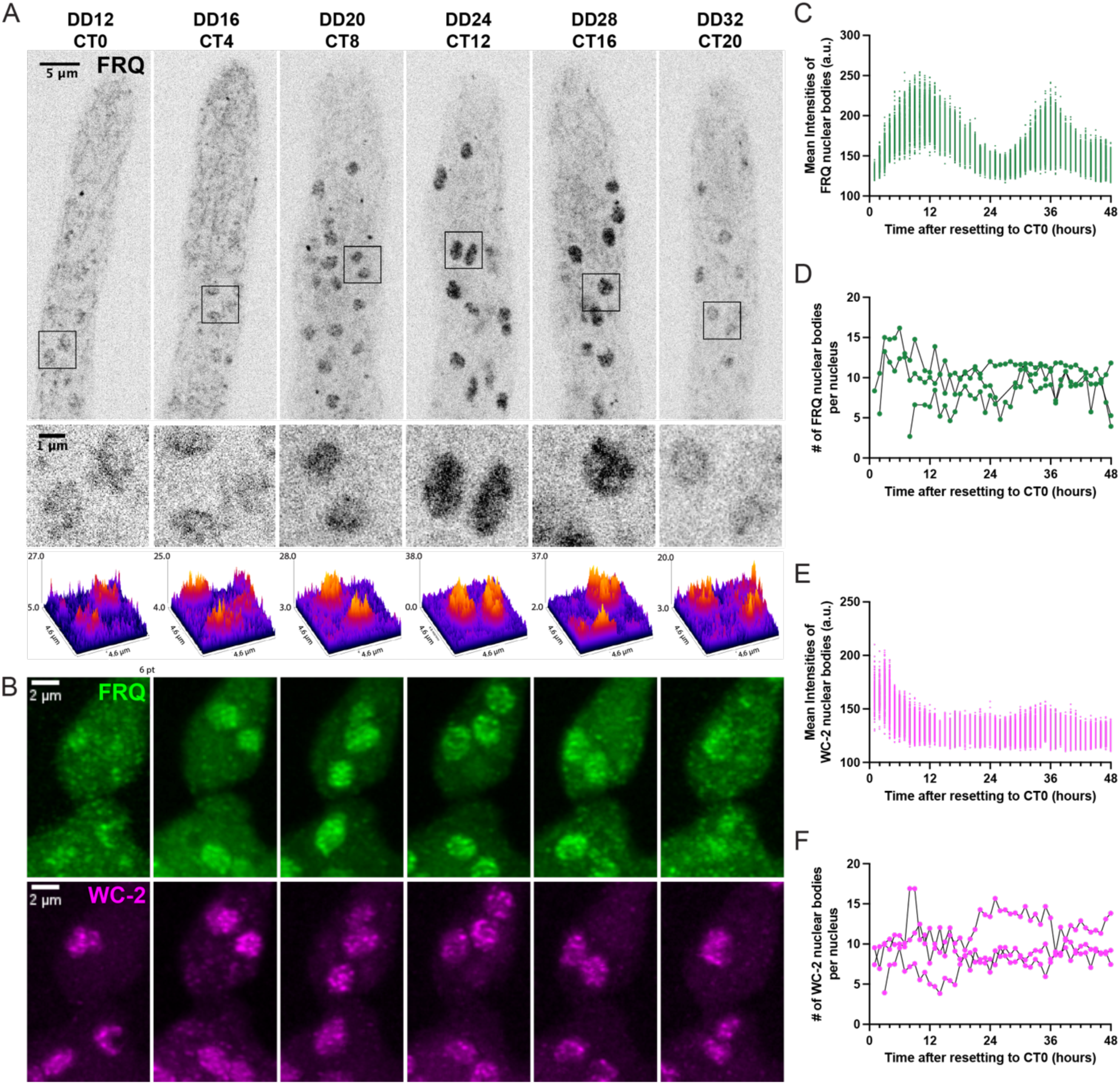
Subnuclear nuclear bodies of clock proteins always exist and remain highly dynamic across circadian cycles. (A) SoRa single focal plane images of FRQ^mNG^ across one circadian cycle. The top panel shows one example of a hyphal tip at each timepoint. The middle panel shows the zoom-in of areas indicated by black squares in the top panel. The bottom panel includes the surface plots of the middle panel, with the y axis as the fluorescent intensity above background (a.u.). (B) Maximal z-projections of 3D deconvolved images of FRQ^mNG^ (top) or WC-2^mAp^ (bottom) at representative timepoints from a 1-hour resolution time course, showing the same circadian timepoints (CT) as in (A). FRQ^mNG^ and WC-2^mAp^ images are contrasted differently for visualization. (C-F) Properties of nuclear bodies identified in 3D deconvolved time courses. n = 3 cells. (C) Fluorescent intensities of FRQ nuclear bodies across circadian cycles. Each dot represents the mean intensity of one nuclear body. (D) Average numbers of FRQ nuclear bodies per nucleus across circadian cycles. Each dot represents the average number calculated for a single cell. Black lines connect values from the same cell across timepoints. (E) Fluorescent intensities of WC-2 nuclear bodies across circadian cycles. (F) Average numbers of WC-2 nuclear bodies per nucleus across circadian cycles.

### Oscillation of FRQ binding correlates the mobility of WCC nuclear bodies

We examined the persistence and mobility of subnuclear bodies as a function of time-of-day to gain insights into their role in timekeeping. Most FRQ nuclear bodies and some WC-1 nuclear bodies were impossible to track reliably due to their high dynamic nature; nonetheless, a subset of WC-1 nuclear bodies could be tracked, and these offer insights into circadian function (Fig. 6A). Using the same stringent parameters (see *Materials and Methods*, “Image analysis”) so as to be sure the same object was tracked between frames, we identified tracks of stationary WC-1 nuclear bodies and saw an oscillation in their average track length/duration (Fig. 6B). Few tracks can be identified for FRQ at any time using the same parameters. Given that we permitted only minimal movement between frames, the track length represents the dwell-time of these stationary WC-1 nuclear bodies. Based on the established nature of transcription factors, we hypothesize that WC-1 specifically bound to target DNA would experience greater physical constraint, leading to a longer dwell time, compared to WC-1 undergoing either free diffusion or non-specific transient binding while searching for targets (Lu and Lionnet, 2021; Jonge et al., 2022; Gebhardt et al., 2013). Our experimental setup excluded freely diffusing WC-1; tracks with a length exceeding five consecutive frames (> 0.5 s) were classified as long-dwell nuclear bodies, indicative of specific binding events, while shorter tracks (≤ 0.5 s) were considered as non-specific binding events. In the subjective morning around CT0-5, WC-1 nuclear bodies on average stay stationary for longer, with larger portion of them being long-dwell nuclear bodies, and their mobility oscillates throughout the circadian cycle (Fig. 6B-C). This is consistent with previous ChIP data indicating that WCC binds rhythmically to the *frq* C-box promoter element (and presumably other C-box-like elements in the genome) in constant darkness, with peak binding at around subjective dawn (CT 20-5) (Belden et al., 2007b; Hurley et al., 2014) (Fig. 6D). Stationary WC-1 nuclear bodies have the shortest dwell time in light, which may be associated with the much higher levels of WCC seen in light, or differences in conformation and DNA binding kinetics between L-WCC and D-WCC (Fig. 6B-C) (He and Liu, 2005; Smith et al., 2010; Hunt et al., 2010; Dasgupta et al., 2015; Wang et al., 2016).

**Figure 6.**
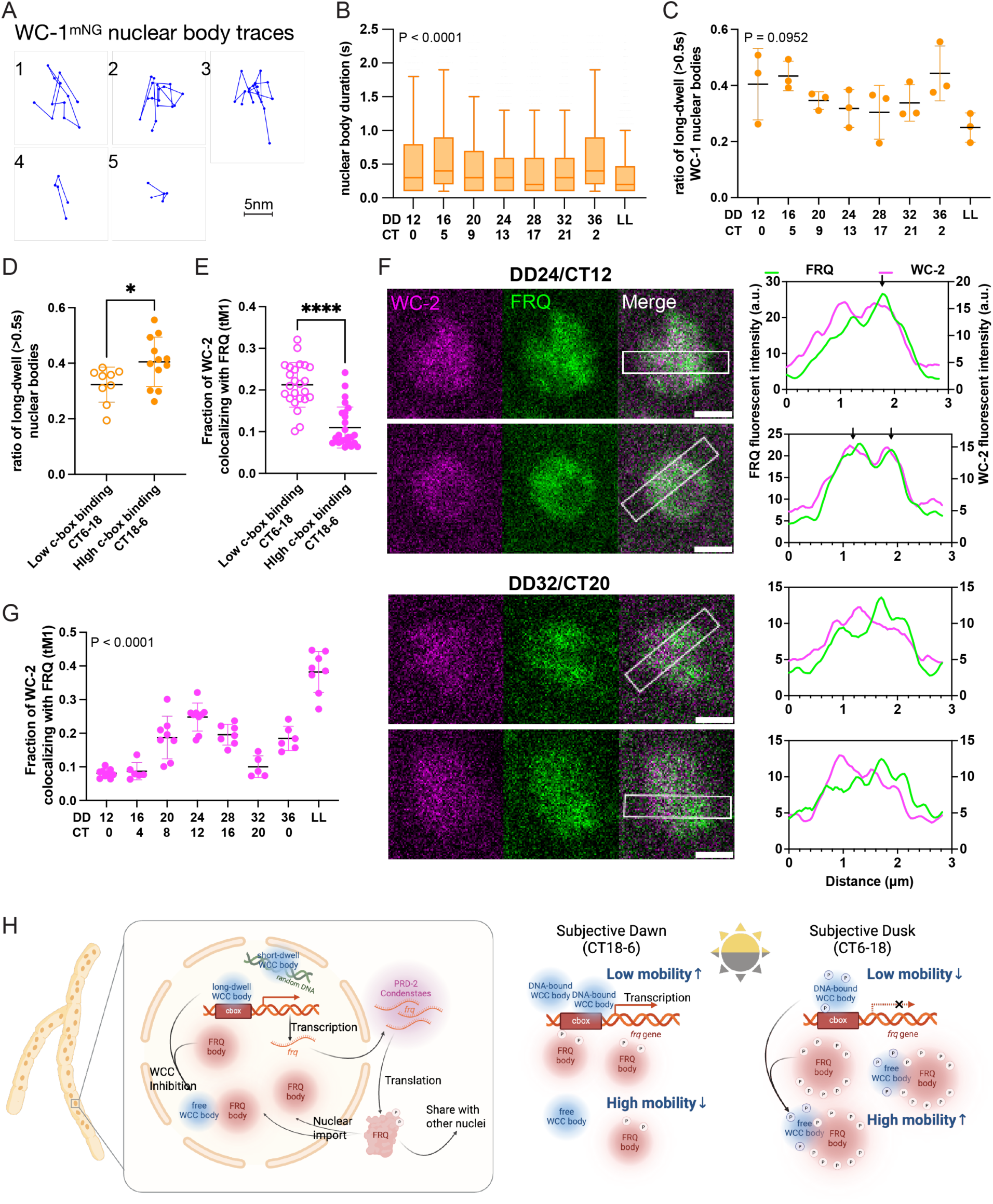
Oscillation of FRQ binding correlates with the mobility of WCC nuclear bodies. (A) Example movement tracks of stationary WC-1 nuclear bodies (dwell time;: 0.2 s). 1-3: long duration tracks (dwell time > 0.5s). 4-5: short duration tracks. (B) Dwell time of stationary WC-1 nuclear bodies across more than one circadian cycle. Tukey-style box plot with median indicated. n = 3-4 hyphae. P < 0.0001 by one-way ANOVA test. (C) Ratio of long-dwell WC-1 versus all stationary WC-1 nuclear bodies across more than one circadian cycle. n = 3-4 hyphae. P = 0.0952 by one-way ANOVA test. (D) Ratio of long-dwell WC-1 nuclear bodies versus all stationary WC-1 nuclear bodies during low *C-box* binding phase (CT 6-18) or high *C-box* binding phase (CT 18-6), n = 3-4 hyphae for every 4 hours. P = 0.0232 by Welch’s t test. (E) Quantification of WC-2 colocalized with FRQ using Manders’ coefficient (tM1) during low or high *C-box* binding phase, N = 5-9 hyphae for every 4 hours. P < 0.0001 by Welch’s t test. (F) Representative dual-camera SoRa images (left) and corresponding profile plots along the white boxes (right) of WC-2 (magenta) and FRQ (green) at example timepoints during either low or high *C-box* binding phases. Black arrows point out overlapping peaks between channels, indicating possible colocalization of nuclear bodies. Scale bar = 1 μm. Images are contrasted differently for visualization. (G) Quantification of WC-2 colocalized with FRQ at each time across more than one circadian cycle using Manders’ coefficient (tM1). P < 0.0001 by one-way ANOVA test. (H) Model illustrating how the circadian clock is regulated at the subcellular and subnuclear levels in a syncytial context, emphasizing on the regulatory role of FRQ binding on WCC mobility and function in circadian clock.

FRQ/FRH-promoted phosphorylation and inhibition of WCC represent a critical step in the TTFL (Dunlap and Loros, 2017; Schafmeier et al., 2005; He et al., 2006; Wang et al., 2019; Lee et al., 2000; Froehlich et al., 2003; Belden et al., 2007b). Previous genetic and biochemical studies indicate this process is mediated by a physical interaction between FRQ and WCC (Cheng et al., 2003; Schafmeier et al., 2005; Baker et al., 2009; Wang et al., 2019; Denault et al., 2001; Cheng et al., 2001; Merrow et al., 2001). Although the formation of nuclear bodies of FRQ and WCC are independent, we wondered if their interactions would regulate the dynamics and thus biological functions. To evaluate their possible interaction *in vivo* across the circadian cycle, we performed co-localization analysis using a dual-camera microscope system, capturing FRQ and WC-2 simultaneously in a high spatial resolution 36-hour time course taken at 4-hour intervals.

Co-localization between WCC and FRQ within nuclei oscillates around a generally low level in the dark, peaking around CT12/DD24 with visually identifiable overlap of some, but never all, FRQ and WC-2 nuclear bodies (Figure 6E-G, Fig. S5I). We were unable to determine if the interaction/co-localization is transient or stable due to difficulties in accurate tracking; specifically, while continuous overlap was observed, it was unclear if these represented the same nuclear bodies across frames due to the extreme mobility of FRQ. The mobility of WCC nuclear bodies correlates with the timing of FRQ-WCC interaction (Fig. 6D-E).

In summary, the changing interaction between FRQ and WCC correlates with their nuclear dynamics and functions in circadian clock. A possible scenario across the day envisions that during the subjective day, FRQ gradually accumulates in nuclei and interacts with WCC, as indicated by increasing co-localization. FRQ then alters WCC function by recruiting kinases, which then phosphorylate WCC (He and Liu, 2005; Schafmeier et al., 2005; He et al., 2006; Wang et al., 2019) while also phosphorylating FRQ itself. Inhibited WCC loses its ability to specifically bind DNA and instead associates with FRQ nuclear bodies having higher dynamics, consequently exhibiting shorter dwell times around subjective dusk (Fig. 6H). As highly phosphorylated FRQ loses its affinity for WCC it is freed to re-bind DNA the next day.

## DISCUSSION

Life is different in a syncytium, and the syncytial organization of *Neurospora* makes it an appealing system for studying cell biology in a multinucleate context. However, live-cell fluorescent imaging in *Neurospora* poses technical challenges. First, the WCC functions as both the primary photoreceptor and as the Positive Arm of the circadian clock feedback loop (Crosthwaite et al., 1997; Talora et al., 1999; He et al., 2002; Cheng et al., 2002), making it difficult to tease apart the effects of laser exposure during fluorescent imaging from intrinsic light/clock regulation. Second, the expression level of core clock components, including FRQ and WCC, is tightly controlled at a low level for proper circadian function (Aronson et al., 1994b; Merrow et al., 1997; Zhou et al., 2013) which limits the achievable signal-to-noise ratio and highlights the need for fluorophores with high brightness and photostability (Wang et al., 2023b). Finally, the dense hyphal network, fast growth rate, and rapid cytoplasmic streaming, combined with the syncytial nature of *Neurospora* hyphae, adds substantial complexity to imaging analysis (Lee et al., 2016; Wang et al., 2024; Bowman, 2025). In this study, we overcame these challenges (Fig.1); our results from continual imaging using microfluidic devices are consistent with existing genetic and biochemical data for FRQ and WCC (Schafmeier et al., 2005; He et al., 2006; Denault et al., 2001; Bartholomai et al., 2022; Hong et al., 2008; Cha et al., 2011; Wang et al., 2014), confirming the validity of the method while providing a rich context of *in vivo* spatiotemporal information. We characterize the spatial and temporal organization of clock components underlying circadian regulation at subcellular and subnuclear scales, revealing how robust circadian rhythms are maintained in complex, dynamic cellular contexts.

Filamentous fungi have complex subcellular organization and require long-distance communication to coordinate their growth and behaviors (Fischer, 1999; Simonin et al., 2012; Fricker et al., 2017; Steinberg et al., 2017; Schmieder et al., 2019; Nagy et al., 2020; Bowman, 2025). Nuclear autonomy is a known feature of syncytial tissues and has been described in filamentous fungi (Bursztajn et al., 1989; Fogarty et al., 2011; Dundon et al., 2016). In *Neurospora*, data suggest that not all nuclei transcribe *frq* transcripts simultaneously (Bartholomai et al., 2022), suggesting a division of labor among nuclei that might increase mRNA usage efficiency (Zinn-Brooks and Roper, 2021). Despite the transcriptional heterogeneity, the distribution of clock proteins appears to be homogenous between nuclei (Fig. 2). These observations, together with prior mRNA data (Bartholomai et al., 2022), are consistent with a model in which some nuclei may contribute *frq* transcripts to maintain the core oscillator while others specialize in circadian outputs with WCC driving clock-controlled genes. Indeed, single-nuclei RNA-seq data in other syncytia has revealed heterogeneous gene expression (Kim et al., 2020; Gerber et al., 2022; Arutyunyan et al., 2023) and other evidence in fungi documenting nuclear differentiation (Jalihal and Gladfelter, 2025) support the concept of nuclear specialization.

The observation that mature *frq* transcripts appear not to freely diffuse in the cytoplasm (Bartholomai et al., 2022) together with the estimated scale of the proposed synchronizing factor (Cheong et al., 2022) and mathematical modeling (Zinn-Brooks and Roper, 2021), imply a requirement for internuclear communication at the protein level. We hypothesize that local sharing of FRQ protein is the key to a harmonious circadian clock and investigated this through a combination of genetic approaches and direct visualization in live cells (Fig.3). Our observation that the clock is insensitive to the proportion of *frq* producing nuclei in heterokaryons aligns with prior results (Loros et al., 1986), as well as with mathematical modeling predicting that a small portion of *frq*-producing nuclei is sufficient to maintain an intact circadian clock in a syncytium (Zinn-Brooks and Roper, 2021) along with buffering transcriptional noise (Little et al., 2013; Mayer et al., 2024). The ability to share WCC components among nuclei potentially adds an additional layer of noise reduction to ensure coherent circadian behaviors.

We also investigated subnuclear dynamics of clock proteins to gain more insights on how spatial organization contributes to circadian regulation. Clock proteins are highly disordered (Pelham et al., 2020; Hurley et al., 2013; Dunlap and Loros, 2018) and have been proposed to undergo liquid-liquid phase separation (LLPS) in multiple model organisms (Conrad et al., 2016; Pelham et al., 2023). Recent *in vitro* experiments demonstrated phase separation ability of FRQ (Tariq et al., 2024). *In vivo*, we identified heterogeneous subnuclear structures of both FRQ and WCC that are much smaller and more dynamic than classic LLPS-like foci (Fig.4), more similar to microbodies of clock proteins recently reported in mammalian cells (Xie et al., 2023). These are unlike the larger and more stable condensates seen in *Drosophila* when tagged proteins are driven by their native promoters (Xiao et al., 2021) or when tagged mammalian or *Neurospora* clock proteins are overexpressed in mammalian cells (Xie et al., 2023; Li et al., 2023; Schunke et al., 2025). While the true nature of the foci, dynamic microbodies vs stable condensates, and the degree to which overexpression may influence observations requires more work, this aspect of regulation is widely seen; nonmembrane-bound foci have been associated with regulation of the clock and circadian output at different levels, including transcription (Xiao et al., 2021; Zhu et al., 2023), post-transcriptional mRNA processing (Bartholomai et al., 2022), translation (Castillo et al., 2022; Zhuang et al., 2023), post-translational modification (Cohen et al., 2014), and environment sensing and response (Jung et al., 2020; Mo et al., 2022; Brown and Nagel, 2025) across model organisms. This common theme across species points to the importance of rapid, reversible protein dynamics in building a stable yet flexible clock.

Strikingly, FRQ forms a stable number of nuclear bodies throughout the circadian cycle (Fig.5) despite robust oscillations in nuclear FRQ abundance and the increasing post-translational modifications, such as phosphorylations, on FRQ across the circadian day that influence its conformation and physical properties (Baker et al., 2009; Tang et al., 2009; Wang et al., 2023a). Unlike the oscillation in the number of nuclear bodies reported in mammal (Xie et al., 2023), or in both number and brightness in *Drosophila* (Xiao et al., 2021), we found that only the brightness of the nuclear bodies of the Negative Arm oscillates in *Neurospora*. Given that the size of nuclear FRQ bodies is close to the diffraction limit, it cannot be directly measured but can be approximated by its brightness. Because condensates maintain a defined concentration inside and outside, the number of molecules within each nuclear body is expected to scale with size and thus with brightness. *In vitro* data suggest that hypophosphorylated FRQ, which is dominant during subjective morning when nuclear FRQ concentration is low, favors LLPS and has a low concentration threshold. As the cycle progresses, FRQ gradually accumulates and gets more phosphorylated, raising both the nuclear concentration and the LLPS threshold, thereby maintaining consistent LLPS capacity (Tariq et al., 2024). Thus, FRQ nuclear bodies might vary in molecular composition and function throughout the circadian cycle without an oscillation in the numbers of FRQ nuclear bodies. The functional importance of these persistent subnuclear foci of the Negative Arm has not yet been fully determined, but they may act as interaction hubs that regulate clock output (Pelham et al., 2023; Coca-Ruiz and Boy-Ruiz, 2025; Costa et al., 2025) and/or buffer concentration to reduce signaling noise (Saunders et al., 2012; Stoeger et al., 2016; Banani et al., 2017; Klosin et al., 2020).

Technical advances enabled this work. Advanced microscopy, including SoRa, provided insights into complex subnuclear organization, though accurate volumetric tracking of FRQ,like PER2, is still unachievable due to technical limitations (Xie et al., 2023). Other techniques used to study molecular dynamics, such as FRAP and FC(C)S, have been applied to clock proteins in mammals (Smyllie et al., 2016, 2022, 2025), but the relatively small size of *Neurospora* nuclei (∼2 µm diameter) together with their rapid and erratic movements makes these techniques particularly challenging. Likewise, microfluidic devices offered a powerful platform for live-cell imaging in 4 dimensions (Ket Chin et al., 2016; Richter et al., 2022; Bedekovic and Brand, 2022). While previous attempts to image circadian output in *Neurospora* over extended periods of time employed different microfluidic designs, overcrowded hyphal growth and (presumably) frequent fusion events made accurate segmentation of single hyphae nearly impossible (Lee et al., 2016; Cheong et al., 2024). Our use of growth mutants (Belden et al., 2007a; Wang et al., 2024) unrelated to clock function and the addition of high–molecular weight polyacrylic acid minimized crowding and enabled continual observation of spatiotemporal control of distributions and dynamics *in vivo* (Wang et al., 2023b) within the same hypha over a circadian timescale (Fig. 1).

Phase separation of Transcription Factors (TFs) is gaining attention as a fundamental mechanism regulating gene expression (Hnisz et al., 2017; Boija et al., 2018; Basu et al., 2020; Chong et al., 2022; Saad et al., 2025; Yang et al., 2024). Phase separation of the Positive Arm in circadian clock has also been proposed and investigated (Pelham et al., 2020; Xiao et al., 2021; Xie et al., 2023; Schunke et al., 2025); however, whether it is intrinsic or dependent on the Negative Arm is debatable. In *Drosophila* clock neurons, dCLK foci only exist during the repression phase and are entirely dependent on dPER (Xiao et al., 2021), whereas in mammals BMAL1 nuclear bodies exist all the time and co-localize rhythmically with PER2 at a low level (∼10%) (Xie et al., 2023); these results are similar to FRQ/WCC in Neurospora (Fig.4), and even at peak most WCC nuclear bodies remain separate from FRQ, in line with biochemical data (Cheng et al., 2005; Schafmeier et al., 2005; Wang et al., 2016; Hong et al., 2008). Across the circadian cycle, WCC mobility changes in line with its promoter binding affinity, with increased residence time during the subjective dawn when *C-box* binding is high (Fig. 6). These timescales align well with established transcription factor (TF) dynamics: freely diffusing TFs explore the 3D nuclear space, briefly scanning accessible DNA (immobile for <1 second), while bound TFs reside on DNA for longer (immobile for 1–100 seconds) (Gebhardt et al., 2013; Lu and Lionnet, 2021; Jonge et al., 2022). Similarly, in mouse neurons, a DNA-binding mutant significantly reduced BMAL residence time, even though the mobility of BMAL shows minimal change between different circadian timepoints by FRAP (Yang et al., 2020; Koch et al., 2022). Dissecting condensate function is complicated by the overlap of intrinsically disordered regions mediating both phase separation and direct binding. Further studies using fast high-resolution imaging and targeted mutagenesis will be essential to clarify these mechanisms.

## MATERIALS AND METHODS

### Strains

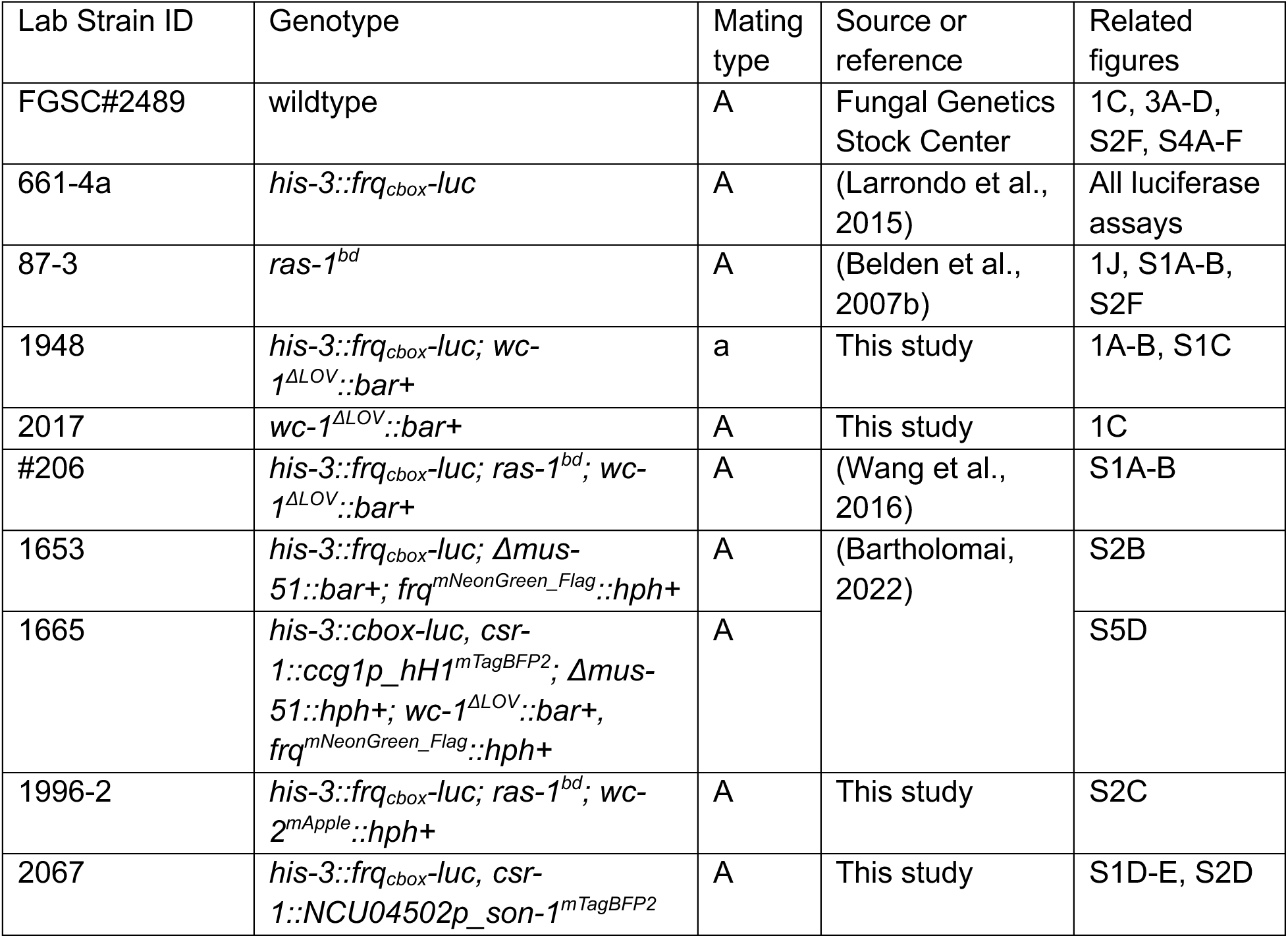
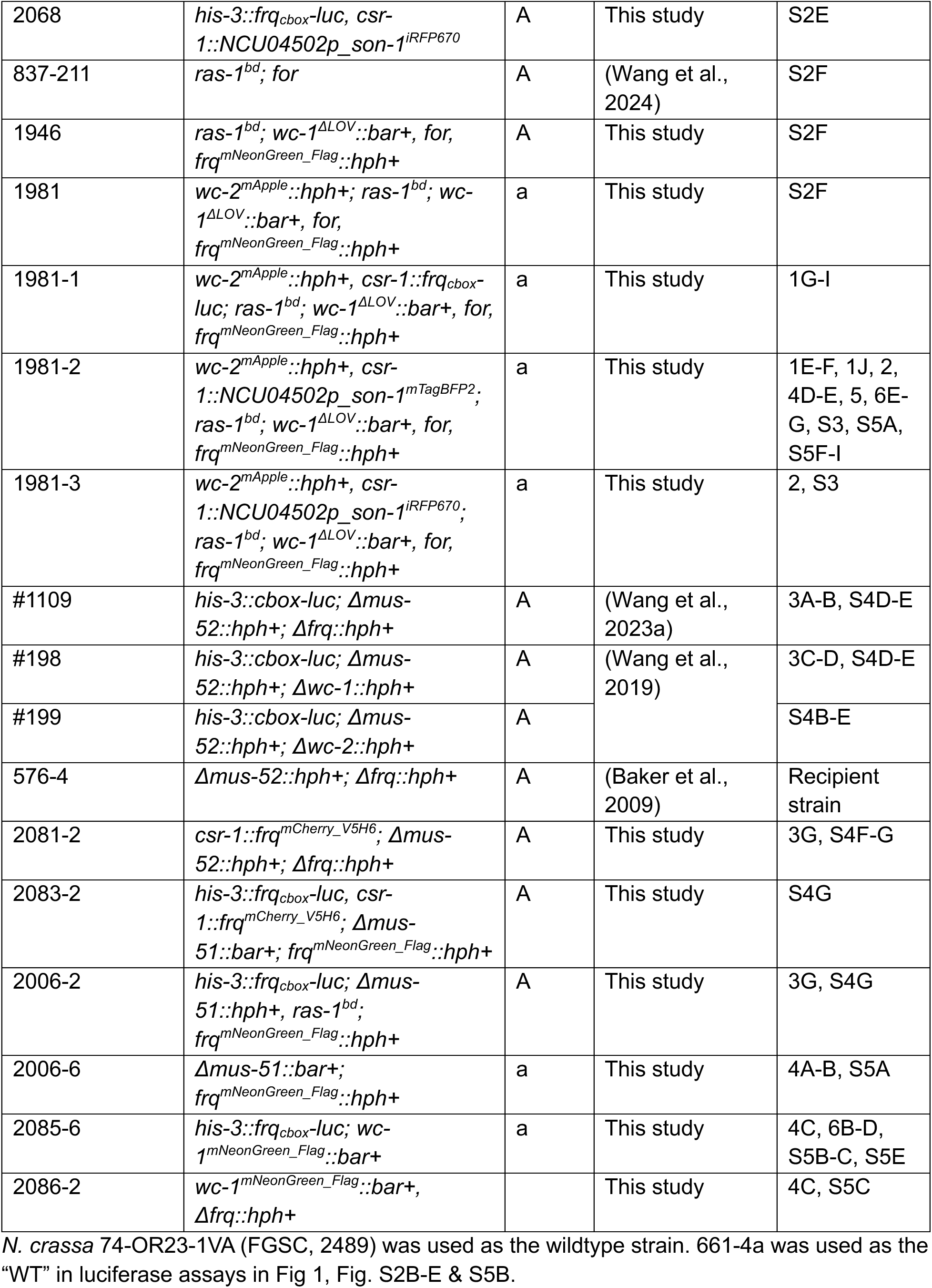
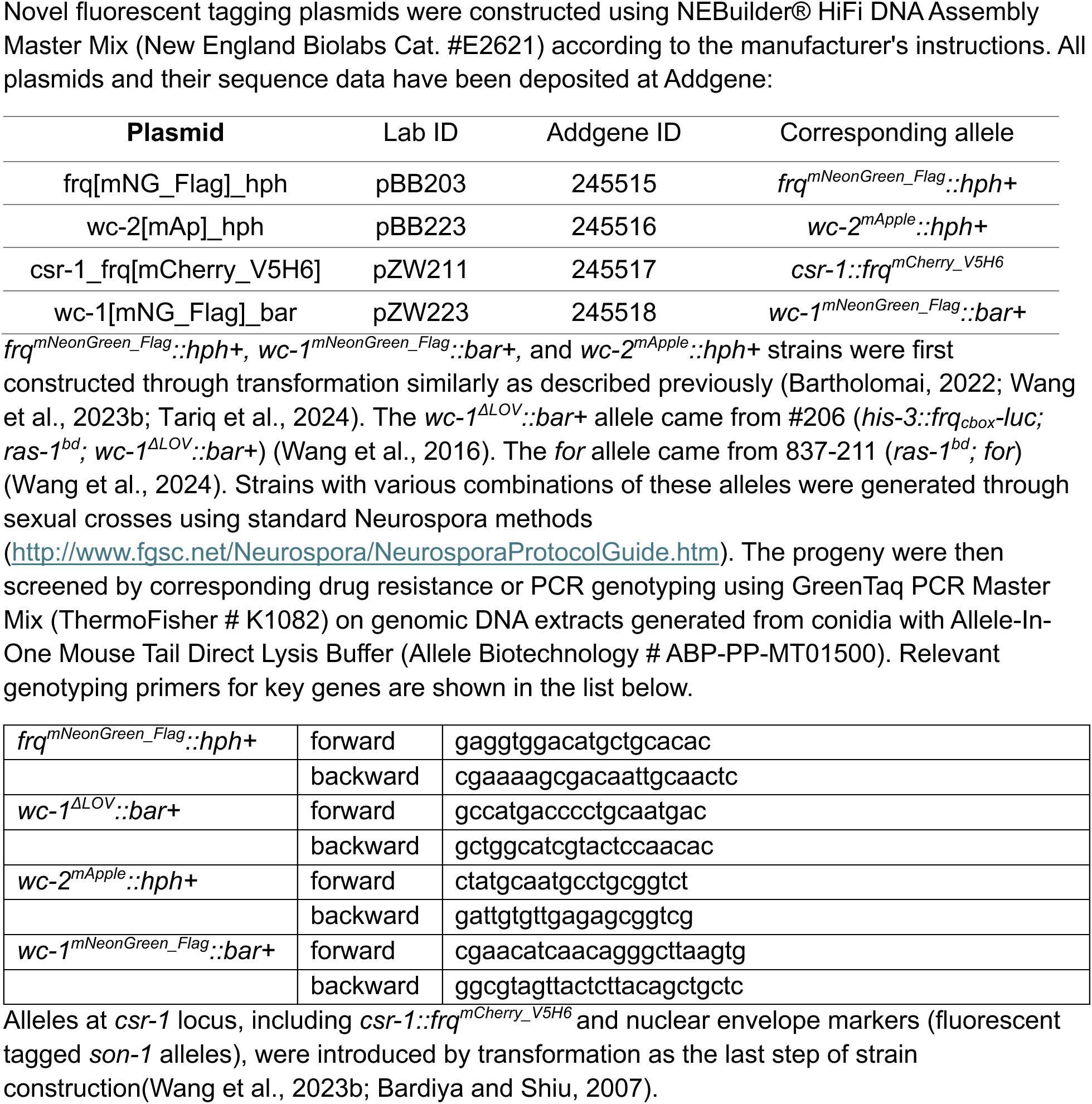

### Luciferase assay

Conidial suspensions were inoculated into black 96-well plates with formate supplemented luciferin medium (1× Vogel’s Salts, 0.1% glucose, 0.17% arginine, 1.5% agar, 50 ng/ml biotin, 25 μM luciferin from GoldBio # 115144-35-9, and 10mM sodium formate from Thermo Scientific #141-53-7), which were then covered by with a Breathe-Easy strip (USAScientific). For luciferase assays generating temperature compensation profiles, inoculated plates were then entrained using 12:12 dark:light cycles at 25°C:20°C, 30°C:25°C, or 33°C:28°C for two days before being transferred to constant darkness at 20°C, 25°C, or 28°C, respectively, for recording. Other luciferase assays were entrained using 2-day 12:12 dark:light cycles at 25°C without temperature changes. Heterokaryons (Figure 3) were made by inoculation of a 1:1 mixture of conidial suspensions from both origin strains, all with the same mating type so they can naturally fuse, unless different conidial inoculation ratios were specified. Recording of luciferase signals was performed by a CCD camera (Pixis 1024B, Princeton Instruments) for 15 minutes every hour for 5 days using LightField software (Princeton Instruments, 64-bit version 6.10.1)(Larrondo et al., 2015). Mean intensities of each well were quantified and background-subtracted in ImageJ using a custom ImageJ Macro(Larrondo et al., 2015). Circadian periods were determined with a custom R program(Kelliher et al., 2020) using data from 12-85 hours post synchronization.

### Race tube assay

Race tube medium (1× Vogel’s Salts, 0.1% glucose, 0.17% arginine, 1.5% agar, 50 ng/ml biotin) supplemented with 10mM sodium formate (Thermo Scientific #141-53-7) or the same medium lacking glucose was poured into long glass race tubes. Conidial suspensions were then inoculated at one end of the race tubes. Growth fronts in each race tube were marked every 24 hours for 5 days as previously described(Dunlap and Loros, 2005; Belden et al., 2007a). The culture conditions are indicated in corresponding figures.

### Protein isolation and detection

Conidia were inoculated into 50mL liquid culture medium (LCM) (1× Vogel’s Salts, 2% glucose, 0.5% arginine, 50 ng/mL biotin) and cultured at 125 rpm in constant light at 25°C for 2 days unless otherwise specified. Cultures were vacuum-dried and then flash frozen in liquid nitrogen. Frozen tissues were hand-ground into fine powders with mortars and pestles and then mixed with protein extraction buffer (50 mM HEPES [pH 7.4], 137 mM NaCl, 10% glycerol, 0.4% Triton X-100) with cOmplete™, Mini, EDTA-free Protease Inhibitor Cocktail (Roche, Catalog # 04693159001). After 10 minutes of incubation on ice, the mixtures were centrifuged at 12,000 rpm for 10 min at 4°C. Protein concentrations in supernatants were determined by Bradford assay, mixed with NuPAGE™ LDS Sample Buffer (ThermoFisher Scientific, Catalog # NP0007) to achieve equal loading per well, and then incubated at 99°C for 5 minutes.

30 μg of total protein were run in each well of a NuPAGE™ 3-8% Tris-Acetate Mini Protein Gels (ThermoFisher Scientific, Catalog #EA03785BOX) in NuPAGE™ Tris-Acetate SDS Running Buffer (ThermoFisher Scientific, Catalog #LA0041). Mini Gel Tank and Blot Module Set (ThermoFisher Scientific, Catalog # NW2000) was used for both electrophoresis and transferring onto PVDF membrane according to the manufacturer’s instructions. PVDF membranes were blocked in SuperBlock T20 (PBS) Blocking Buffer (ThermoFisher Scientific, Catalog # 37516) for 30 minutes at room temperature before antibody incubations.

Immunoblotting was carried out in PBS buffer containing 0.2% Tween-20 (PBS-T): primary antibodies were incubated overnight at 4°C, secondary antibodies were incubated for 1 hour at room temperature, with 3 PBS-T washes between steps. The chemiluminescence signals were developed with Pierce™ ECL Western Blotting Substrate (ThermoFisher Scientific, Catalog #32106), then detected using Azure c400 imaging system (Azure Biosystems) and analyzed in FIJI.

| Antibodies | Source | Dilution |
| --- | --- | --- |
| Anti-FRQ (rabbit) | (Hurley et al., 2013) | 1:250 |
| Anti-WC-1 (rabbit) | (Froehlich et al., 2002) | 1:250 |
| Anti-WC-2 (rabbit) | (Denault et al., 2001) | 1:2000 |
| Anti-αTubulin (mouse) | Santa Cruz DM1A | 1:5000 |
| Anti-V5 (mouse) | Invitrogen #46-0705 | 1:5000 |
| Goat Anti-Rabbit IgG (H + L)-HRP Conjugate | BIO-RAD #1706515 | 1:5000 |
| Goat Anti-Mouse IgG (H + L)-HRP Conjugate | BIO-RAD #1706516 | 1:5000 |

### Microfluidic device fabrication

The device for cell trapping consists of a flow layer of 25 μm height and cell layer of 8 μm height. The device to monitor cell fusion has a single layer of 8 μm height. The master molds were fabricated by ultraviolet photolithography using standard methods. SU-8 (Microchem) photoresist was applied to a silicon wafer by spin coating to appropriate thickness (corresponding to the channel height) and patterns were then created by exposing the uncured photoresist to ultraviolet light through film masks (CAD/Art Services).

Dimethyl siloxane monomer (Sylgard 184) with 1:20 curing agent was mixed and added on top of the wafer mold, and air bubbles were removed using a vacuum chamber. After baking at 65°C overnight, holes were introduced using a biopsy punch to connect the feeding channels to the external tubing, and individual chips were cut and bonded onto KOH-cleaned cover slips using oxygen plasma treatment. After plasma treatment, the individual chips were immediately put on an 80°C hotplate for 30 min and post-baked at 65°C overnight.

### Growth in microfluidic device

Conidia are taken with a wooden stick from 6/7 days old complete medium slants and put into 1ml sterile water. Conidia suspensions were filtered with 10μm syringe filter, and then washed with another 1ml sterile water. For inoculation, filtered conidia suspensions were injected using a 1ml syringe with a 5/8’’ blunt end needle until liquid fills up the chamber. After waiting for 20 minutes for conidia to distribute throughout the channels, a syringe pump (New Era Pump Systems, NE-300 Just Infusion™ Syringe Pump) with a 10ml syringe and tubing was used to propel media through the device at a constant rate of 1.5 μl/min.

For 2-day continuous recordings (Figure 2), Bird medium(Metzenberg, 2004) with 0.01% glucose, 0.2% polyacrylic acid and 1mM formate was used. After inoculation, the culture was kept at 30°C for 10h and then transferred to 20°C for 6h for entrainment before imaging at 26°C. The temperature is maintained by Tokai Hit Stage Top Incubator during imaging. Two individual runs with sibling strains 1981-2 and 1981-3 were conducted.

For merging/fusion experiments (Figure 3), inoculation inlets and outlets were sealed by small pieces of metal wire after co-inoculation of conidia suspensions of 2006-2 (*frq^mNeonGreen^*) and 2081-2 (*frq^mCherry^*) in opposite inoculation channels. Fresh LCM was then flowed through the media channel and the culture was kept and imaged at room temperature.

### Spinning disk confocal microscopy

All fluorescent images were acquired on one of two Nikon Eclipse Ti-E stand-based spinning disk confocal microscope systems with Nikon LU-N4 laser launch that includes 405 nm, 488 nm, 561 nm, and 640 nm lasers. One of the systems has a Yokogawa CSU-W1 spinning disk system, and the other is the Nikon SoRa system with a 2.8X magnifier and a 4.0X magnifier. Both systems are equipped with 2 Photometrics Prime BSI sCMOS cameras, an ASI MS-2000 motorized stage with a piezo Z-drive as well as a Tokai Hit stage-top incubation system. All images were acquired with a Nikon CFI Plan Apochromat λD 60× oil objective with a numerical aperture of 1.42. All experiments described in this section were performed using the conventional spinning disk mode. Experiments conducted with the SoRa mode are described separately in a later section.

For recording of growth in whole microfluidic devices (Fig.1E, S1D-E, Supplemental Video 1&3), extra-large single focal images were obtained by stitching multiple Fields of View (FOVs) together in NIS-Elements every hour.

For 2-day continuous recordings (Fig. 2), multiple xy positions were imaged sequentially every hour for > 48 hours on the CSU-W1 system. A dual-or tri-color 10 μm z-stack of 35 slices with 300 nm step size was taken sequentially for each position and each time point.

### Image analysis

All 3D segmentation was performed using a customed pipeline in Arivis Vision4D.

For 2-day continuous recordings (Fig. 2), segmentation of hypha and nuclei was based on red channel (WC-2^mApple^) using the same parameters for all timepoints. Only nuclei within the target hyphal object were recognized and analyzed. Each cell ended up with one hyphal object, tens of nuclear objects, and a manually generated background object for each timepoint. Information including the mean, sum fluorescent intensity, and volume of each object was exported for analysis. 23 cells from 2 individual runs were analyzed.

For nuclear signal quantifications (Fig. 2B), the combined mean FRQ fluorescent intensity of all nuclear objects for each time point and each cell was calculated using the following equation:

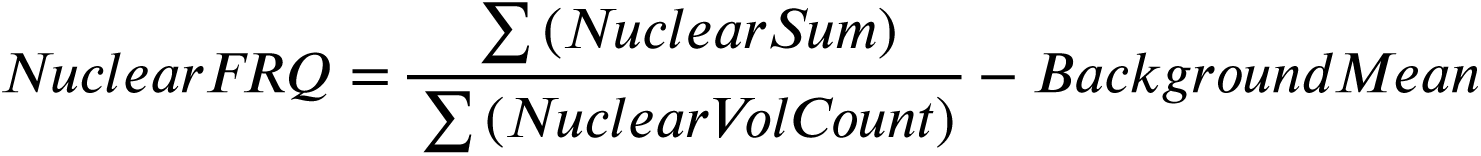

*NuclearSum* is the sum fluorescent intensity of each nuclear object; *NuclearVolCount* is the volume of each nuclear object in arbitrary units. *BackgroundMean* is the mean intensity of the background at the corresponding timepoint. The same calculation was done for WC-2 (Fig. 2C). All other data plotted in Figure 2 were subtracted by the corresponding *BackgroundMean* as well.

3D deconvolution of 2-day continuous recordings (Fig. 5B) was performed in Nikon Elements software using the Richardson–Lucy algorithm with 11 iterations, followed by 3D segmentation of nuclear bodies in Arivis Vision4D using Blob Finder. The same parameters (Diameter = 0.324μm; Possibility threshold = 20%; Split sensitivity = 20%) were used for all timepoints and only objects overlapping with previously identified nuclear objects were included for analysis. Segmentations for FRQ nuclear bodies and WC-2 nuclear bodies were performed separately. Mean intensity of each nuclear body object and the number of objects identified at each timepoint were exported for analysis. If the number of nuclear bodies per nucleus was smaller than 3 at a given timepoint, it was considered as invalid segmentation (failed to identify nuclear bodies correctly using the given parameters by eye) and the data is excluded. 3 cells were deconvolved and analyzed.

### SoRa imaging and analysis

7 samples at 4-hour resolution in constant darkness and 1 sample in constant light for each strain were prepared for rhythmic timeseries as previously described(Loros et al., 1989). All samples were grown on agarose pads (1× Vogel’s Salts, 2% sucrose, 0.5% arginine, 50 ng/mL biotin, 1% agarose) and then imaged using the “inverted agar block” method(Hickey et al., 2004).

All timelapse imaging of hyphal tips at growing fronts was performed with 100ms exposure time and no delay between frames for up to 5 seconds (50 frames) using the SoRa mode with 2.8X magnifier and 60x objective. Single channel SoRa images for FRQ^mNG^ or WC-1^mNG^ were performed using the single-camera model with 488nm laser for excitation and 515 – 555 nm for emission detection (Fig. 4A-C, 5A, 6A-D, and S5A&E, Supplemental Video 5). Dual-camera model was used for 2-channel SoRa images (Fig. 4D and 6F, Supplemental Video 4) to ensure simultaneous Green/Red imaging considering the high dynamic behavior. 488nm and 561nm lasers were used for excitation. The built-in dichroic mirror (DM A561LP) directs light of wavelength > 561 nm to Camera 1 as the red channel (shown in magenta) and reflects the rest to Camera 2 as the green channel.

The movements of WC-1^mNG^ nuclear bodies were analyzed using TrackMate in FIJI(Ershov et al., 2022). 5-second timelapses were first corrected for photobleaching. In TrackMate, objects were identified using the DoG detector (optimal for small spot sizes) with the estimated object diameter as 0.4 μm. Built-in simple LAP tracker with stringent parameters (linking max distance: 0.2 μm, gap-closing max distance: 0.2 μm, gap-closing max frame gap: 1) was used for tracking. Therefore, only stationary tracks (500-700 tracks per recording) were identified. The same parameters were used for all timepoints. For each time point, 3 recordings with least nuclear movements were included in the analysis. Each recording/FOV included 1-2 hyphal tips with tens of nuclei.

For colocalization between FRQ and WCC, the dual-camera images were background subtracted (background = 100 for both channels) and then analyzed using JACoP in FIJI(Bolte and Cordelières, 2006). The red and green channel thresholds were set to 15 and 12, respectively, at all timepoints to detect only bright spots. For each time point, 5-9 FOVs were included in the analysis. Each FOV included 1-2 hyphal tips with tens of nuclei. Data for profile plots were extracted in FIJI using a 2.15 μm × 0.6 μm ROI (Fig. 4D) or a 2.85 μm × 0.6 μm ROI (Fig. 6F), then smoothed (5 neighbors averaged, 4th-order smoothing polynomial) and plotted in Prism 10.

### Statistical analysis and data visualization

Statistical analysis data visualization was performed using Prism 10 (GraphPad). All statistical tests and the number of replicates (n) used for each experiment are indicated in the relevant figure legends. All schematic diagrams were created in BioRender.

## ACKNOWLEDGMENTS

This study was supported by a National Institutes of Health (NIH) grant awarded to Jay C. Dunlap (R35GM118021). Daniel Schultz was supported by NSF grant PHY 2412766. David Ritz was supported by the Dartmouth PhD Innovation Fellowship program. We acknowledge use of stocks from the Fungal Genetics Stock Center (FGSC.net), NIAID-supported informatic resources at FungiDB (Contract HHSN75N93019C00077), and Molecular Biology facilities supported by the Dartmouth CQB COBRE (NIH NIGMS grant P20GM130454), and the Dartmouth BioMT (NIH NIGMS grant P20-GM113132). Imaging was performed in the Life Sciences Light Microscopy Facility (RRID:SCR_027143) at Dartmouth College, with support from NIH award to Yashi Ahmed (S10OD032310) for purchase of the super-resolution spinning disk confocal.

## Notes

### Competing Interest Statement

The authors have declared no competing interest.

