## Supplemental Materials for "Core Clock Protein Subcellular Dynamics Coordinate Local and Global Circadian Control in Syncytia"

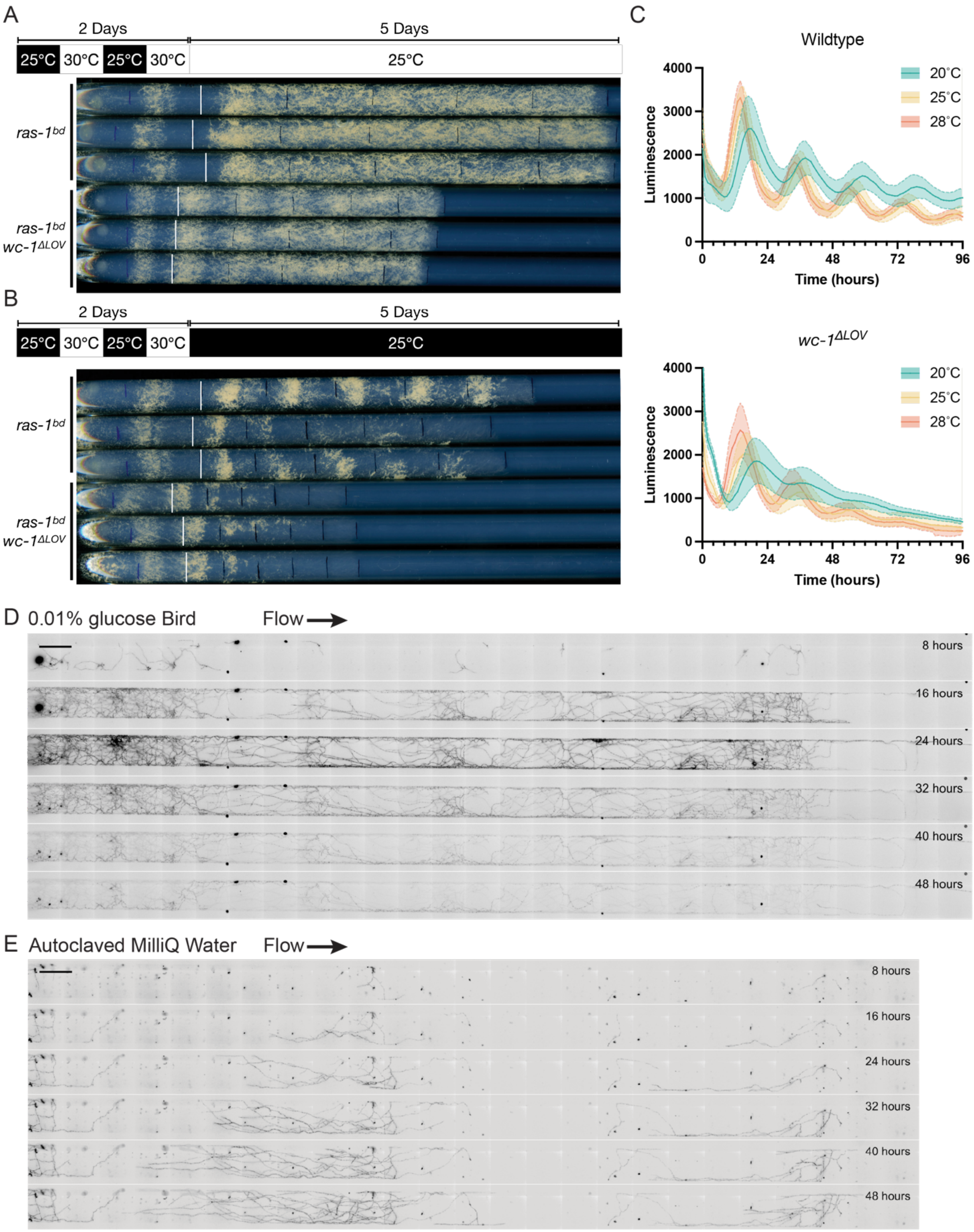
A
2 Days
5 Days
25°C 30°C 25°C 30°C
25°C
ras-1bd
ras-1bd
wc-1ΔLOV
B
2 Days
5 Days
25°C 30°C 25°C 30°C
25°C
ras-1bd
ras-1bd
wc-1ΔLOV
C
Wildtype
20°C
25°C
28°C
Luminescence
Time (hours)
wc-1ΔLOV
20°C
25°C
28°C
Luminescence
Time (hours)
D 0.01% glucose Bird
Flow
8 hours
16 hours
24 hours
32 hours
40 hours
48 hours
E Autoclaved MilliQ Water
Flow
8 hours
16 hours
24 hours
32 hours
40 hours
48 hours

Figure S1. Growth features of live *Neurospora* with blind *wc-1*$^{\Delta LOV}$ mutation or in customized microfluidic devices.

(A-B) Race tube assays of clock wildtype and *wc-1*$^{\Delta LOV}$ strains in (A) constant light or (B) constant darkness. The schematic diagram shows the entrainment and culture conditions. The strains were entrained by 12h:12h light and temperature cycles before being released to constant conditions. (C) Luciferase assays of clock wildtype (strain 661-4a, top) and *wc-1*$^{\Delta LOV}$ (strain 1948, bottom) across temperatures. Each curve represents the average of three biological triplicates each with 4 technical replicates, with the shade showing standard deviations. (D-E) Example timelapse showing the growth of wildtype strain with nuclear marker *csr-1:: son-1*$^{mTagBFP2}$ (strain 2067) in the microfluidic device with fresh flow of (D) medium or (E) water. Images were taken in DAPI channel monitoring the fluorescent signal of nuclear envelope marker SON-1$^{\text{mTagBFP2}}$. Scale bar = 200µm.

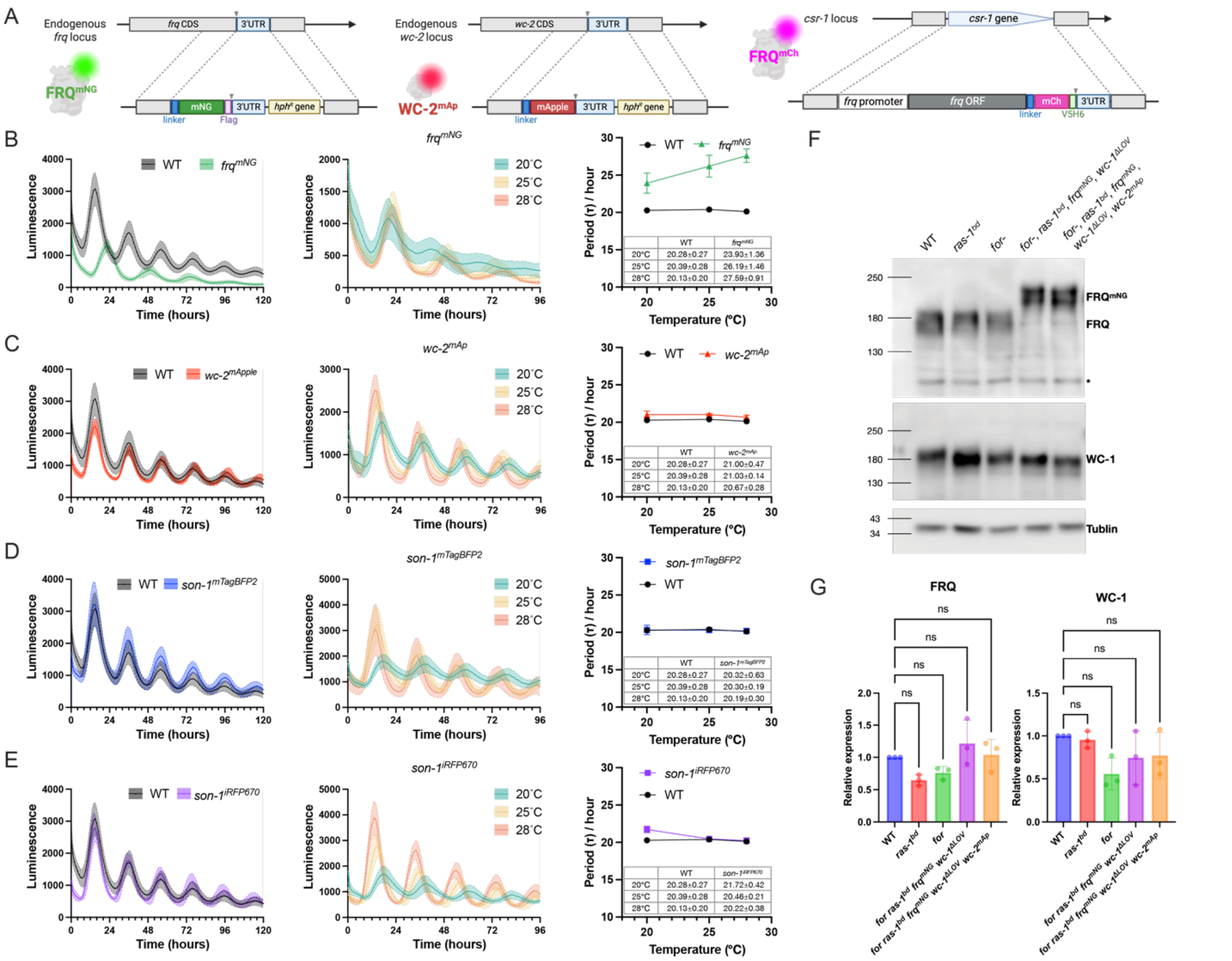


Figure S2. The circadian clock retains function when core clock proteins are tagged by various fluorescent protein markers.

(A) Schematics of tagging strategies whereby clock proteins are tagged with markers at their endogenous loci or at a neutral location using homologous recombination. (B-E) Luciferase assays of strains with each single tag: (B) $frq^{mNG}$ (C) $wc\text{-}2^{mAp}$ (D) *csr-1:: son-1*$^{mTagBFP2}$ (E) *csr-1:: son-1*$^{iRFP670}$. The left panels are luciferase traces of the corresponding strain at 25℃ compared to wildtype; the middle panels are luciferase traces of the corresponding strain across temperatures. The right panels are periods calculated from luciferase assays at different temperatures after temperature entrainments compared to wildtype. n=12 replicates for each condition. Error bars show one standard deviation. Full genotypes are in the strain table in Materials and Methods. (F) Western blots showing expression levels of core clock proteins in imaging strains with fluorescent tags. (G) Quantification of core clock protein expression levels from 3 biological replicates. Statistical comparisons were performed using one-way ANOVA followed by Dunnett's multiple comparisons test.

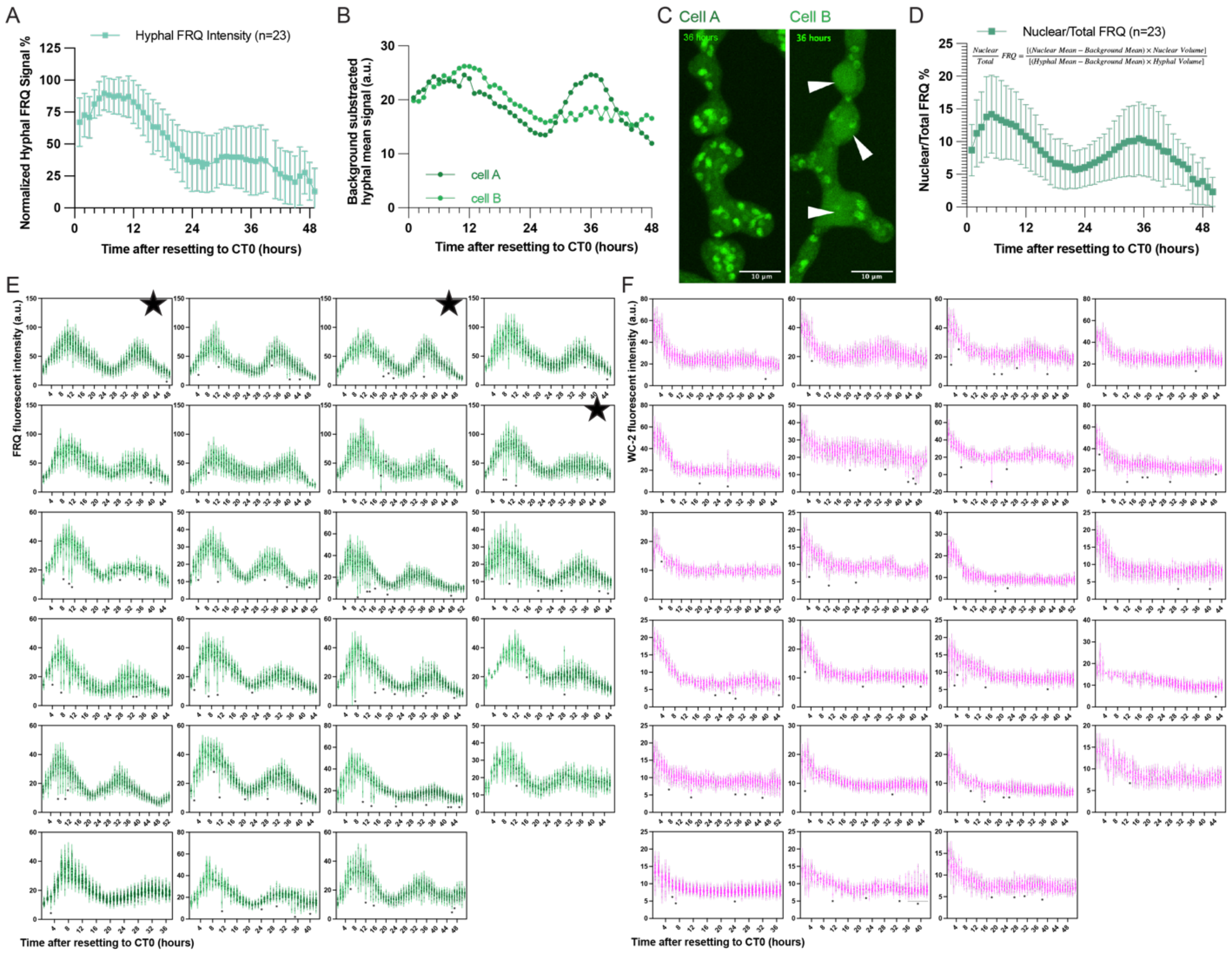


Figure S3. FRQ and WC-2 translocate to all nuclei across circadian cycles.

(A) Average fluorescent intensities of FRQ in whole hyphae. n=23 cells. The maximal and minimal signals for each cell are set as 100% and 0% for normalization. All error bars show one standard deviation. (B) Whole hyphal FRQ intensities of 2 representative cells shown in (C). Cell A represents cells without vesicles thereby allowing accurate quantification of total FRQ across time. Cell B, in contrast, accumulated large vesicles in the second cycle, pointed out by white arrowheads, which interfere with accurate quantification of total FRQ. (D) The proportion of nuclear FRQ molecules among total FRQ calculated using the indicated equation. n=23 cells. (E) Distributions of FRQ signal in each nucleus over time for every cell. 3 cells marked by the black stars were used for deconvolution analysis in Figure 5. (F) Distributions of WC-2 signal in each nucleus over time for every cell. Figure 2D-E are representative examples of (E-F).

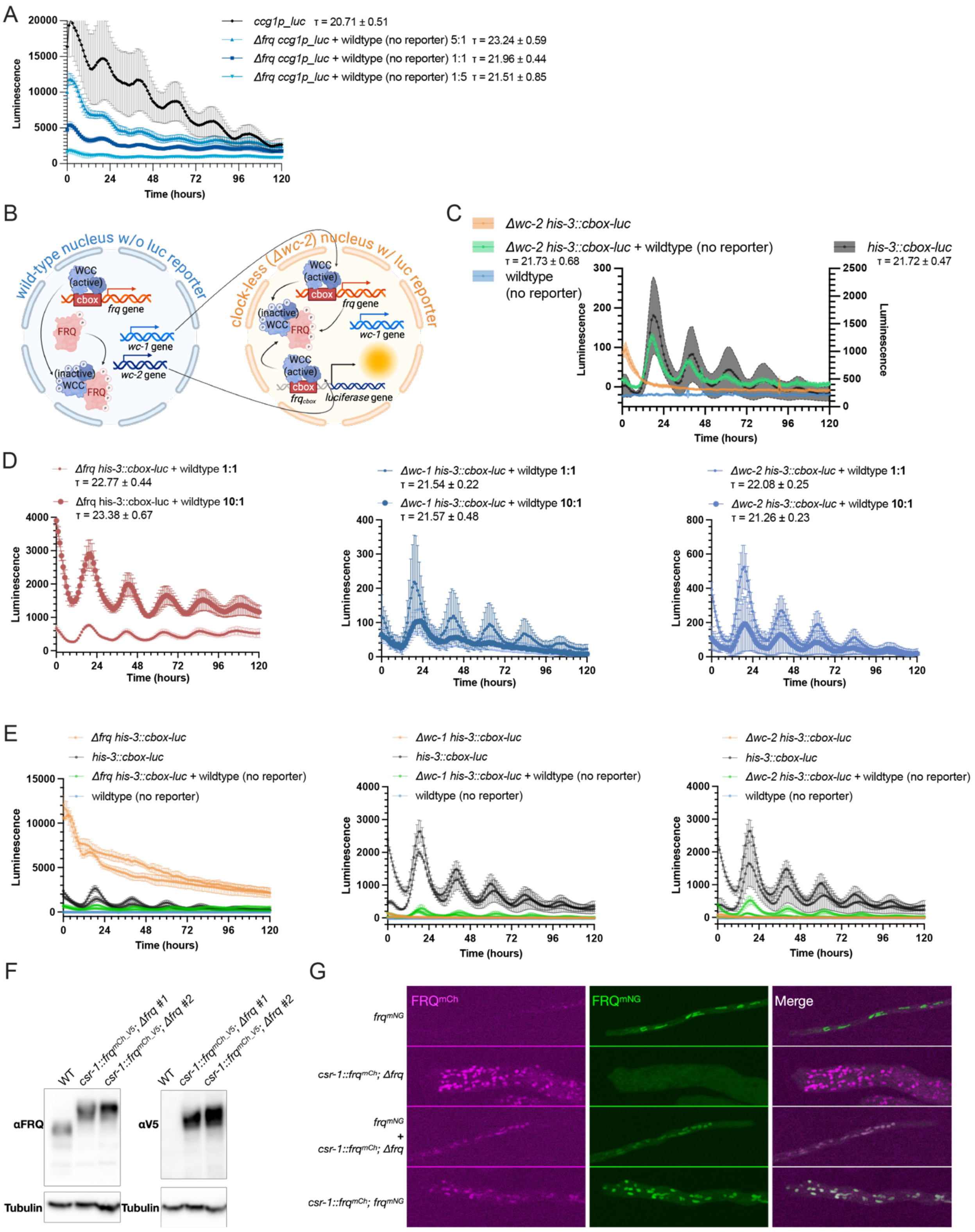
A
ccg1p_luc τ = 20.71 ± 0.51
Δfrq ccg1p_luc + wildtype (no reporter) 5:1 τ = 23.24 ± 0.59
Δfrq ccg1p_luc + wildtype (no reporter) 1:1 τ = 21.96 ± 0.44
Δfrq ccg1p_luc + wildtype (no reporter) 1:5 τ = 21.51 ± 0.85
Luminescence
Time (hours)
B
wild-type nucleus w/o luc reporter
clock-less (Δwc-2) nucleus w/ luc reporter
WCC (active)
cbox
frq gene
FRQ
wc-1 gene
wc-2 gene
(inactive) WCC
luciferase gene
C
Δwc-2 his-3::cbox-luc
Δwc-2 his-3::cbox-luc + wildtype (no reporter)
τ = 21.73 ± 0.68
wildtype (no reporter)
his-3::cbox-luc
τ = 21.72 ± 0.47
Luminescence
Time (hours)
D
Δfrq his-3::cbox-luc + wildtype 1:1
τ = 22.77 ± 0.44
Δfrq his-3::cbox-luc + wildtype 10:1
τ = 23.38 ± 0.67
Δwc-1 his-3::cbox-luc + wildtype 1:1
τ = 21.54 ± 0.22
Δwc-1 his-3::cbox-luc + wildtype 10:1
τ = 21.57 ± 0.48
Δwc-2 his-3::cbox-luc + wildtype 1:1
τ = 22.08 ± 0.25
Δwc-2 his-3::cbox-luc + wildtype 10:1
τ = 21.26 ± 0.23
Luminescence
Time (hours)
E
Δfrq his-3::cbox-luc
his-3::cbox-luc
Δfrq his-3::cbox-luc + wildtype (no reporter)
wildtype (no reporter)
Δwc-1 his-3::cbox-luc
Δwc-1 his-3::cbox-luc + wildtype (no reporter)
Δwc-2 his-3::cbox-luc
Δwc-2 his-3::cbox-luc + wildtype (no reporter)
Luminescence
Time (hours)
F
WT
csr-1::frqmCh_V5; Δfrq #1
csr-1::frqmCh_V5; Δfrq #2
αFRQ
αV5
Tubulin
G
FRQmCh
FRQmNG
Merge
frqmNG
csr-1::frqmCh; Δfrq
frqmNG + csr-1::frqmCh; Δfrq
csr-1::frqmCh; frqmNG

Figure S4. Local sharing of clock components facilitates circadian control.

(A) Luc traces representing *ccg-1* rhythmicity, as a proxy for all clock output, indicate restoration of circadian output by local sharing of FRQ. Each curve represents the average of 4 technical replicates, with error bars showing one standard deviation. Conidial inoculation ratios and period lengths are marked in the legends. (B) Schematics showing the model for WC-2 local sharing in a heterokaryon of *Δwc-2 his-3::cbox-luc* and wildtype. (C) Luc traces of *Δwc-2 his-3::cbox-luc*, wildtype strains, their heterokaryon, and the positive control strain *his-3::cbox-luc*. The scale for *his-3::cbox-luc* is plotted on the right y axis, and scales for the other 3 traces are plotted on the left y axis, as indicated above the axes. Each curve represents the average of 4 technical replicates with the shade showing one standard deviation. The periods ($\tau$) of rhythmic traces are marked on the figures. A plot with all traces, as well as another set of biological replicates, plotted on the same scale is included in Fig. S4E. (D) Luc traces for heterokaryons with different conidial inoculation ratios. Large dots represent more clock-less nuclei with reporter and correspondingly less clock component; small dots represent more corresponding clock component, less clock-less nuclei with reporter. (E) Data from Fig. 4B, 4D, and Sup. Fig. 4B and their replicates but plotted on the same y scale to show the differences in intensity. (F) Expression level of FRQ in *csr-1:: frq*$^{mCh\_V5}$*; Δfrq* strains compared to wildtype. (G) Representative maximal projections of z-stacks acquired from *csr-1:: frq*$^{mCh}$*; Δfrq* and *frq*$^{mNG}$ strains (used in Figure 4E), their heterokaryon, and a *csr-1:: frq*$^{mCh}$*; frq*$^{mNG}$ strain.

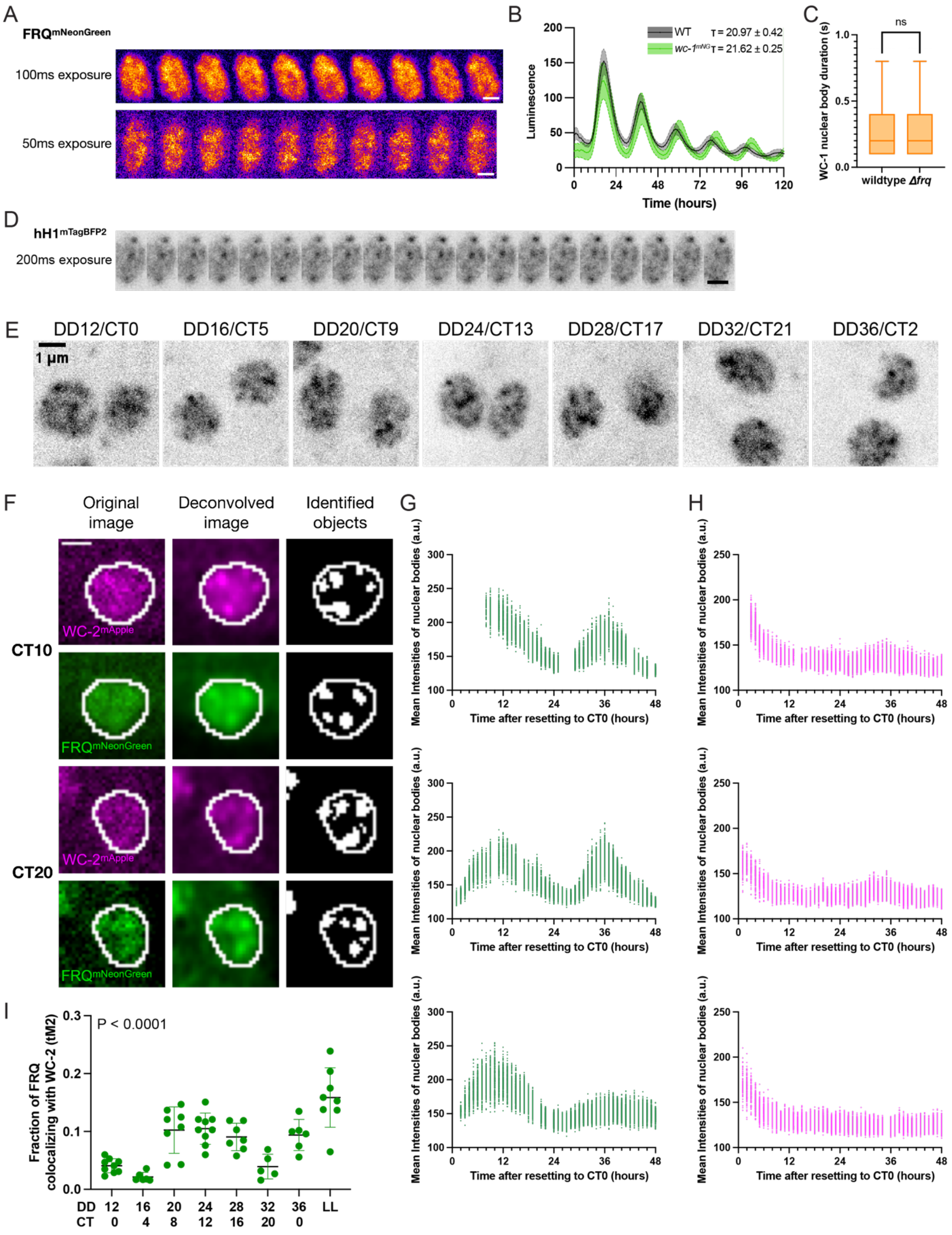

A
FRQ mNeonGreen
100ms exposure
50ms exposure
B
Luminescence
Time (hours)
WT τ = 20.97 ± 0.42
wc-1 mNG τ = 21.62 ± 0.25
C
ns
WC-1 nuclear body duration (s)
wildtype
Δfrq
D
hH1 mTagBFP2
200ms exposure
E
DD12/CT0
DD16/CT5
DD20/CT9
DD24/CT13
DD28/CT17
DD32/CT21
DD36/CT2
1 μm
F
Original image
Deconvolved image
Identified objects
CT10
CT20
WC-2 mApple
FRQ mNeonGreen
G
H
Mean Intensities of nuclear bodies (a.u.)
Time after resetting to CT0 (hours)
I
P < 0.0001
Fraction of FRQ colocalizing with WC-2 (tM2)
DD 12 16 20 24 28 32 36 LL
CT 0 4 8 12 16 20 0

Figure S5. Highly dynamic nuclear bodies of Positive and Negative arm clock components persist across circadian cycles with oscillating colocalization.

(A) Timelapses of $FRQ^{mNG}$ with 100ms (top) or 50ms (bottom) exposure time without delay acquired on the SoRa system. Scale bar = 1μm. (B) Luciferase assays of clock wildtype and *wc-1*$^{mNeonGreen}$ strains at 25℃. Each curve represents the average of biological triplicates each with 4 technical replicates, with the shade showing one standard deviation. Periods (τ). calculated from these traces are marked in the figure. (C) Dwell time of stationary WC-1 nuclear bodies in wildtype or *Δfrq* background. Tukey-style box plot with median indicated. n = 3-4 hyphae. (D) Timelapses of $hH1^{mTagBFP2}$ with 200ms exposure time without delay acquired on the SoRa system. Scale bar = 1μm. (E) SoRa single focal plane images of WC-$1^{mNG}$ across more than one circadian cycle. 2 nuclei of one hyphal tip at each timepoint are shown as examples. Scale bar = 1μm. (F) Middle focal plane images of $FRQ^{mNG}$ (green) or WC-$2^{mAp}$ (magenta) from two example timepoints in the 1-hour resolution time course showing the results of deconvolution and segmentation. The left panel is the original image. The middle panel is the middle focal plane image after 3D deconvolution. The right panel is the binary segmentation result with identified objects shown in white. White circular outlines shared by all 3 panels are nuclear boundaries. Scale bar = 1 μm. (G-H) Fluorescent intensities of (G) FRQ or (H) WC-2 nuclear bodies across circadian cycles of each replicate from Fig. 5C or E, respectively. Each dot represents the mean intensity of one nuclear body. (I) Quantification of FRQ colocalized with WC-2 at each time across more than one circadian cycle using Manders' coefficient (tM2). P < 0.0001 by one-way ANOVA test.

Supplemental Video 1. *N. crassa* grow against the flow independent of nutrient gradients.

Supplemental Video 2. Localization of FRQ and WC-2 across circadian cycles.

Supplemental Video 3. Hyphal fusions can be monitored in microfluidic devices.

Supplemental Video 4. Subcellular dynamics of nuclei and core clock proteins during cytoplasmic streaming and septal pores crossing.

Supplemental Video 5. Positive and Negative Arm clock components form highly dynamic nuclear bodies.
